# Unlocking the therapeutic potential of the corneal endothelium: Intravital imaging reveals endogenous regenerative capabilities

**DOI:** 10.1101/2022.12.15.520611

**Authors:** Maxwell Marshall, Raneesh Ramarapu, Lisa Ohman, Bakary Samasa, Tyler Papp, Parhiz Hamideh, Sangwan Park, Brian C. Leonard, Sara M. Thomasy, Panteleimon Rompolas

## Abstract

Loss of vision due to corneal endothelial dysfunction affects millions worldwide. The development of new treatments is hampered by the incomplete knowledge of the regenerative capacity of corneal endothelial cells *in vivo*. Herein, we developed a mouse model to directly monitor corneal endothelial regeneration in real time, and at the single cell level, by two-photon microscopy. We show that the mouse corneal endothelium recapitulates the main features of human endothelial physiology, including complete cellular quiescence and a decline in cell density with aging. Critically, we demonstrate the endogenous regenerative potential of the tissue by capturing the proliferation of corneal endothelial cells during repair of large injuries. By single cell lineage tracing analysis, we provide evidence that corneal endothelial cells are equipotent in their ability to activate the cell cycle and contribute to tissue regeneration. Based on these findings we developed a feasible therapeutic approach to stimulate the regeneration of the corneal endothelium, using modified mRNA technology. To reprogram corneal endothelial cells in vivo and unlock their ability to escape quiescence, we combined five modified mRNAs encoding for proteins involved in cell cycle activation. Injection of the encapsulated mRNAs directly into the eye of older mice induced transient proliferation of corneal endothelial cells that led to an increase in endothelial cell density, effectively reversing the effect of aging. This therapeutic strategy offers a compelling paradigm for treating ocular disease and modulating tissue regeneration in organs with limited endogenous ability.

## Main Text

A major goal of Regenerative Medicine is to rejuvenate organs whose function is afflicted by aging or disease. The corneal endothelium is critical for healthy vision but has limited documented regenerative activity [1]. The cornea, a clear tissue located at the front of the eye, serves as the primary window of our visual system. Corneal endothelial cells are important for regulating the optical transparency of the cornea by maintaining the collagenous stroma in a state of relative dehydration, through their ion pumping activity and barrier function [2, 3]. These cells are developmentally derived from the neural crest and establish a tissue monolayer in the posterior corneal surface. This monolayer becomes mitotically inactive postnatally and into adulthood, through well studied mechanisms of contact inhibition and insulation from growth signals [4–6]. A progressive decline in the number of corneal endothelial cells, due to aging or other confounding factors, can lead to corneal edema and loss of vision due to a decompensated ion pumping capacity [7]. Corneal endothelial disorders are a leading cause of reduced vision and blindness, affecting ∼4% of the population over 40 [8]. The current standard of care for treating corneal endothelial dysfunction is transplant surgery (penetrating or endothelial keratoplasty), which presents a challenge due to the complexity and cost of the procedure, and worldwide shortages in donor tissue [9, 10].

Despite exciting new prospects, the pace of research to develop alternative therapies for corneal endothelial dysfunction is hampered by the lack of comprehensive knowledge of regenerative capacity of corneal endothelial cells due to the inherent inefficiencies of current experimental models [11]. To address this challenge, we harnessed cutting-edge genetic lineage tracing approaches combined with two-photon microscopy [12] and developed an *in vivo* model to directly visualize regeneration of the mouse corneal endothelium, at the single cell level, by longitudinal live imaging. Our system combines modified mRNA technology to label corneal endothelial cells *in vivo* to track their activity and analyze the direct outcomes of experimental interventions in real time. Proliferation of corneal endothelial cells, in humans as well as other species, has been demonstrated to occur under certain conditions, such as *ex vivo* or in cell culture [13–15]. However, the activity of corneal endothelial cells *in vivo* is still poorly understood. In contrast to large animal models, the mouse is readily amenable to genetic manipulation. Furthermore, mouse knockout models of loci associated with Fuchs’ endothelial corneal dystrophy show clinical features of the disease [16, 17]. We therefore sought to capitalize on contemporary genetic tools to investigate different facets of corneal endothelial physiology, by live imaging. The small size of the mouse eye is advantageous for our system, which can rapidly image the whole cornea and capture cellular behaviors across the entire organ. Thus, we directly imaged the eyes of live mice, using genetically encoded fluorescent reporters to resolve the fine morphology of the corneal endothelium at a single cell resolution (**Fig 1A**).

**Figure 1.**
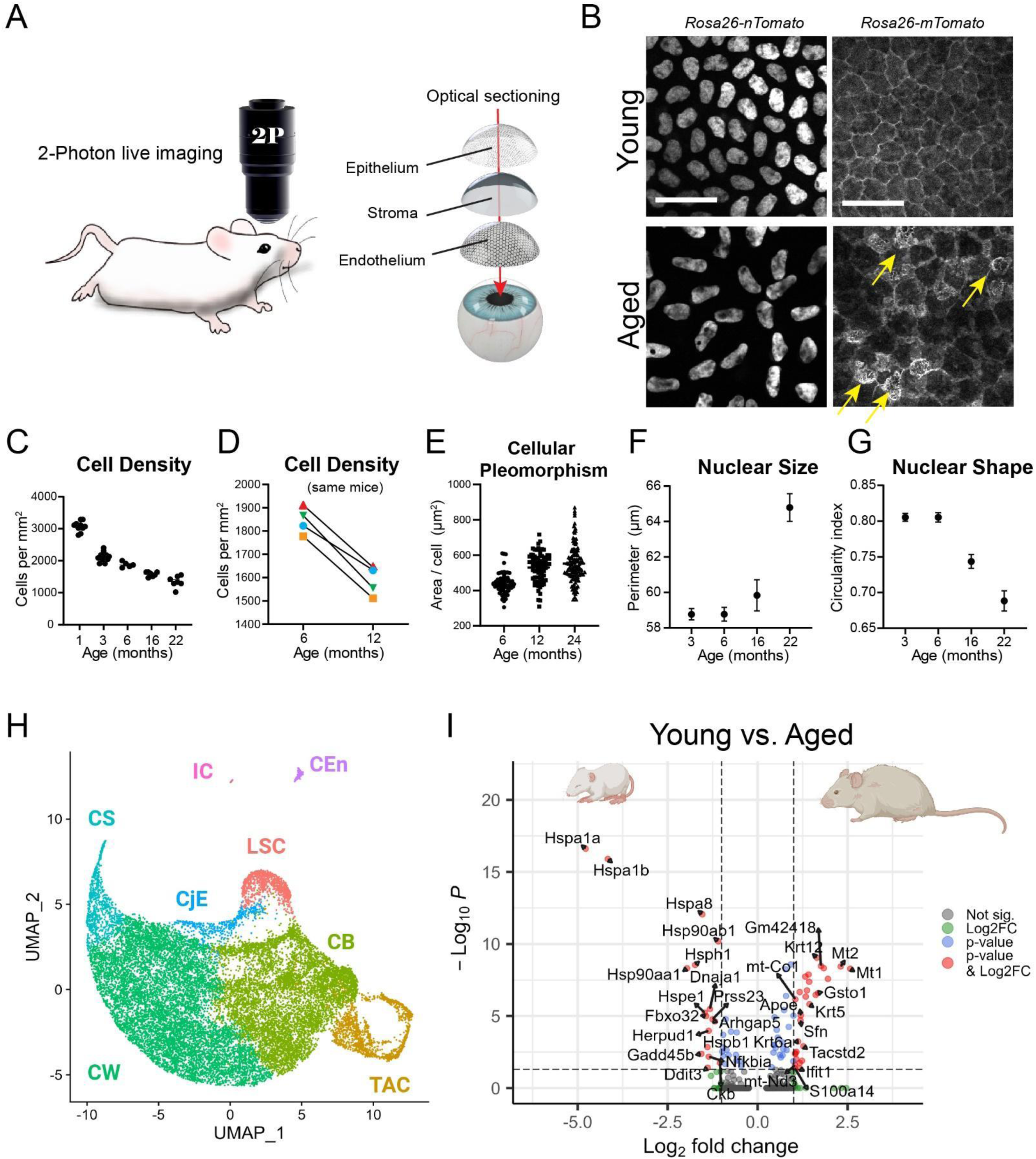
Age-related morphological and gene expression changes in the murine corneal endothelium. A) Experimental approach to directly analyze the physiology and regeneration of the mouse corneal endothelium at the single-cell level, by intravital imaging. Schematic shows optical sectioning of the live mouse corneal endothelium, imaged by two-photon microscopy. Yellow arrows indicate guttae-like structures between corneal endothelial cells. B) High magnification images of the mouse corneal endothelium from 3-month and 22-month-old mice, using membrane- or nuclear-localized in vivo fluorescent reporters. C) Quantification of cell density in the corneal endothelium as a function of age [N = 40, 8 corneas per age group, P < 0.0001; one-way ANOVA]. D) Quantification of cell density in the same corneas imaged at the indicated ages [N = 4 corneas, P < 0.0024; two-way ANOVA]. E) Quantification of cellular pleomorphism in the corneal endothelium as a function of age [N = 254 cells from 6 corneas per age group, P < 0.0001; one-way ANOVA]. Scale bar 50 μm. F) Quantification of nuclear perimeter of corneal endothelial cells as a function of age [N = 241 nuclei from 8 corneas, P < 0.0001; one-way ANOVA]. G) Quantification of nuclear circularity [(4π (Area)/(Perimeter^2^)] of corneal endothelial cells as a function of age [N = 241 nuclei from 8 corneas, P < 0.0001; one-way ANOVA]. H) UMAP plot of the aging scRNA-seq atlas of the mouse cornea. Unbiased clustering of integrated samples across three ages (2, 6, and 11-month-old) in wild type mice revealed 8 cell populations of the cornea (N = 9, 3 sets of corneas per age group). LSC, limbal stem cell; TAC, transient amplifying cells; CB, corneal basal cells; CW, corneal wing cells; CS, corneal superficial; CjE, conjunctival epithelium; CEn, corneal endothelium; IC, immune cells. I) Volcano plot demonstrating upregulated and downregulated genes in the Young CEn (2MO, n = 38) with respect to Aged CEn (6MO + 11MO, n = 99) with 1842 significant DE genes (P < 0.05).

The morphological characteristics of mouse corneal endothelial cells recapitulate main features of the human tissue, including a hexagonal apical surface and ruffled basal cell membranes (**Fig S1**). Common clinical presentations of corneal endothelial dystrophies include: 1) a gradual cell loss, and corresponding changes in the shape and size of individual cells with age, 2) the appearance of guttae-like lesions; extracellular outgrowths that interrupt the lateral contacts between endothelial cells [18, 19]. Live imaging of mouse corneas during different stages of adulthood, showed a consistent cell loss, accompanied by aberrant cellular morphology (polymegathism and pleomorphism), that correlated with age (**Fig 1B-E**). Another unique feature of aged corneas revealed by this analysis, was a pronounced change in nuclear shape and size (**Fig 1B, F-G**), which may indicate variations in chromatin organization and gene expression, as previously proposed for other cell types [20]. These data validated the relevance and sensitivity of our approach and demonstrated that the physiology of corneal endothelial cells can be accurately studied by live imaging.

The observed decline in cell density of the mouse corneal endothelium as a function of aging suggests that corneal endothelial cells do not self-renew under normal homeostasis, at least at a sufficient level to replenish physiological cell loss due to tissue attrition. To further investigate the changes that occur at the molecular level in the corneal endothelium during aging we performed single-cell transcriptional analysis of mouse corneas, harvested from 2-, 6- and 11-month-old mice. Unbiased clustering of the integrated samples across the three ages revealed all the predicted corneal cell populations, based on established gene markers, including a unique cell cluster that corresponds to the corneal endothelium (**Fig 2H, S2A**). Differential gene expression analysis of the identified corneal endothelial cells between young and aged mice revealed changes in key genes (**Fig 2I, S2B**). Specifically, several unique gene markers related to corneal endothelial cell function, such as *Atpb1*, *Col8a2*, *Col4a1*, and *Slc4a11*, were all downregulated in aged mice. Conversely, corneal endothelial cells from aged corneas upregulated keratin genes, including *Krt5*, and *Krt12*, which are normally not associated with corneal endothelial cell identity. Taken together these experiments revealed a decline of the corneal endothelium during physiological aging, which is associated with changes in gene expression and results in progressive cell loss.

**Figure 2.**
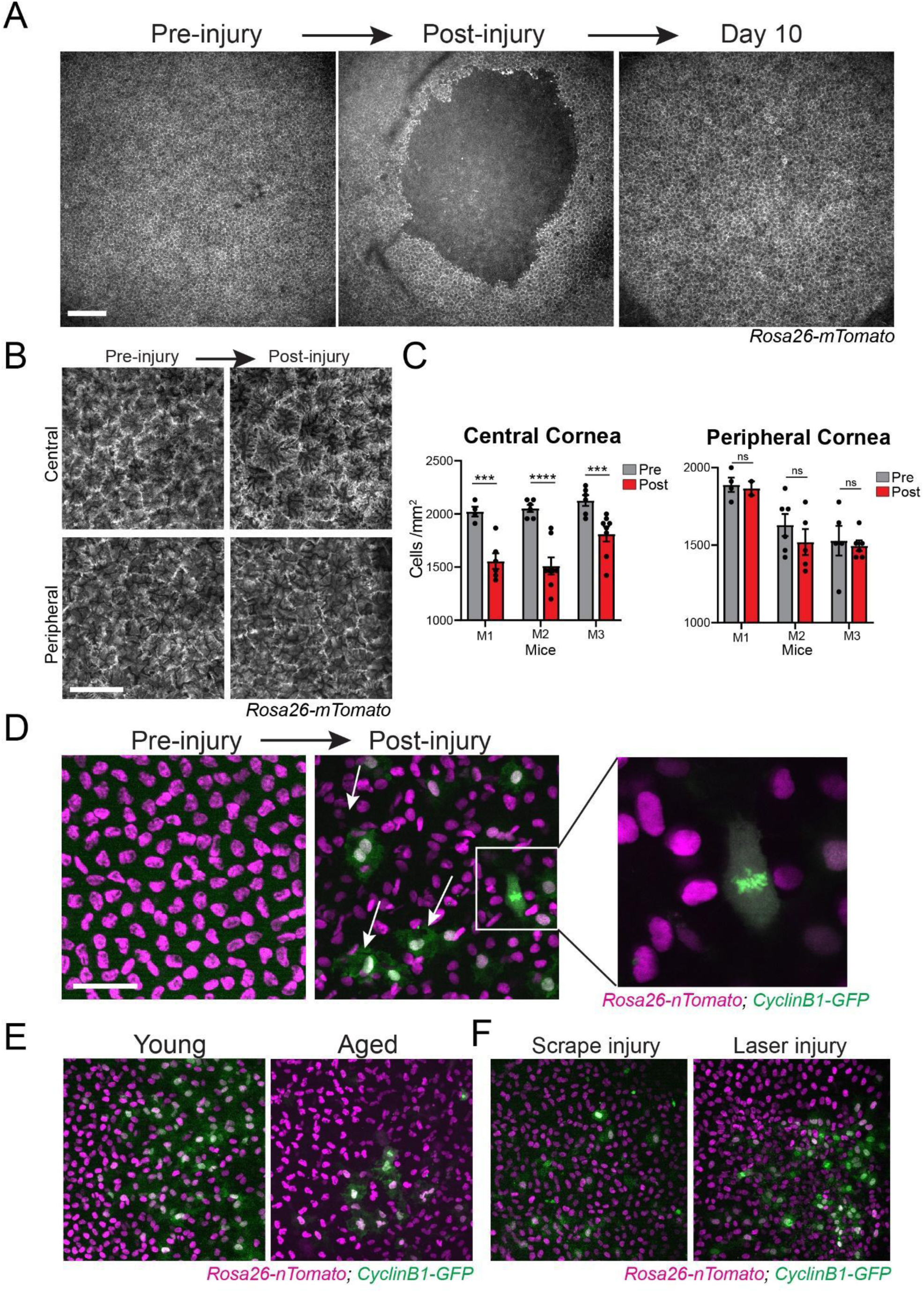
Injury captures the regenerative potential of the corneal endothelium. A) Representative time points from injury repair of the corneal endothelium captured by longitudinal live imaging. Panels show en face view of the corneal endothelium generated by maximum projection of serial optical sections. B) Representative high magnification views from the same eye of the central and peripheral corneal endothelium, imaged before and after injury. C) Quantification of cell density in the central and peripheral regions of the corneal epithelium. Each bar pair represents a different cornea analyzed before and after injury [N = 3-8 fields/cornea, P <0.0015 / 0.0001 / 0.0052 / 0.78 / 0.34/ 0.73, paired t-test]. D) Representative high magnification views of the corneal endothelium taken before and two days after injury. A globally expressed red fluorescent reporter with nuclear localization (*Rosa26-nTomato*) is used to visualize all corneal endothelial cells in the tissue. Actively proliferating cells (arrows) are visualized by green fluorescence from expression of the in vivo cell cycle reporter (*CyclinB1-GFP*). E) Examples of cell proliferation in the corneal endothelium, two days after scrape injury, in Young (3-month old) and Aged (15-month old) mice. E) Examples of cell proliferation in the corneal endothelium, two days after scrape or laser injury. Scale bars 200 μm (A), 50 μm (B-E).

Clinical evidence of partial regeneration after stripping of the corneal endothelium, are consistent with earlier studies in animal models, which suggest that corneal endothelial cells have mitotic potential that can be activated by injury [21–24]. While a mechanism for injury repair of the corneal endothelium is generally accepted, the cellular activities involved in this process and the distinct contribution of individual cells to the regeneration of the tissue have not been fully resolved. Elucidating the conditions that can activate the cell cycle *in vivo* is critical for informing effective therapeutic strategies aiming at induced proliferation in the corneal endothelium. To address this, we developed an *in vivo* injury model to induce localized damage to corneal endothelium and analyze the cellular behaviors during the repair process, by longitudinal live imaging. We found that non-penetrating, superficial scraping of the epithelium causes an immediate rupture of the endothelium in the posterior corneal surface; likely due to acute stromal edema from the loss of epithelial barrier (**Fig 2A, S3**). By re-imaging the same corneas over time, we captured the dynamics of the injury repair across the entire corneal tissue. Within 2-3 days, the corneal endothelium was able to repair the injury, re-establishing a seemingly intact monolayer of cells (**Fig 2A, S3**). However, detailed analysis revealed that the cell density was significantly reduced in the corneal endothelial layer compared to pre-injury levels (**Fig 2B, C, S3**). Moreover, individual corneal endothelial cells were found to be enlarged at various degrees (**Fig 2B**).

The observed aberrant phenotype was more pronounced in the center of the tissue, near the site of injury, suggesting a repair mechanism driven primarily by cell migration and enlargement at the wound edge. Alternatively - or in concert - cell proliferation may be limited to the periphery of the corneal endothelium, where a population with stem cell properties is proposed to reside [25–26]. To directly answer this question, we used mice that express a cell-cycle reporter (CyclinB1-GFP) to capture proliferating cells during the tissue repair, by live imaging. We validated the specificity and faithful kinetics of the reporter by imaging the mitotically active corneal <u>epi</u>thelium (**Fig S4A-B**). Pre-injury, GFP was not detected in the corneal stroma or the <u>endo</u>thelium. Proliferating cells appeared in the corneal endothelium 24 hours post-injury. Their numbers peaked at 2 days and proliferation became again undetectable in the tissue three days after injury (**Fig 2D, S5C-E**). This is consistent with the timing of the repair process and the re-establishment of the corneal endothelial monolayer. Aged mice harboring the cell cycle reporter, showed a similar number of GFP+ cells in the corneal endothelium after injury, compared to young mice (**Fig 2E**). Furthermore, activation and proliferation of corneal endothelial cells was observed as part of the wound healing process in other injury models, including femtosecond laser ablation (**Fig 2F, S4F-H**), suggesting that this response is universal. These data directly demonstrate a proliferative potential of corneal endothelial cells *in vivo* that is maintained with age. More importantly, these experiments show that under these conditions, proliferation is induced in cells located exclusively near the wound margins. This evidence argue against a spatially restricted stem cell population and support the hypothesis that all cells in the corneal endothelium possess regenerative potential. Furthermore, response to local environmental cues appears to be critical for cell cycle activation after injury.

To effectively test this hypothesis, we sought to improve upon the sensitivity of our *in vivo* assays to track the activity of single corneal endothelial cells and analyze their contribution to tissue regeneration. The gold standard in Regenerative Medicine research is the use of Cre/LoxP-based in vivo lineage tracing systems with cell-type specific genetic drivers [27]. To date, such approaches have not been widely implemented to study the regeneration of the corneal endothelium, primarily due to the lack of validated endothelial-specific Cre drivers. To address this, we devised a strategy to induce loxP genomic recombination using Cre-encoding modified mRNA (**Fig 3A**). When delivered systemically, using lipid nanoparticle vehicles, mRNAs are endocytosed primarily by hepatocytes, but localized induced expression has been previously demonstrated when mRNAs are injected in situ [28]. We hypothesized that local delivery of modified mRNA would be especially advantageous for the corneal endothelium, due to the protected environment of the anterior eye chamber and the relatively slow turnover of the aqueous humor. In principle, these conditions would aid the retention and endocytosis of the mRNA cargo by corneal endothelial cells. To test this, we performed direct (intracameral) injection of GFP-encoding mRNA in the eyes of wild type mice and validated expression by live imaging (**Fig 3A, S5**). We observed robust GFP signal in approximately 5-20% of corneal endothelial cells, which peaked 24 hours after injection and persisted for up to 7 days after injection (**Fig 3B, S5**). Elsewhere in the anterior eye segment, expression was detected only in the iris but was completely absent from the corneal epithelium and stroma, or the lens (**Fig S5**). Expression was also not observed in the non-injected contralateral eye (**Fig S5**). Taken together, we show that topical delivery of encapsulated mRNA into the anterior eye chamber can effectively target the corneal endothelium, with likely low toxicity due to off-target effects.

**Figure 3.**
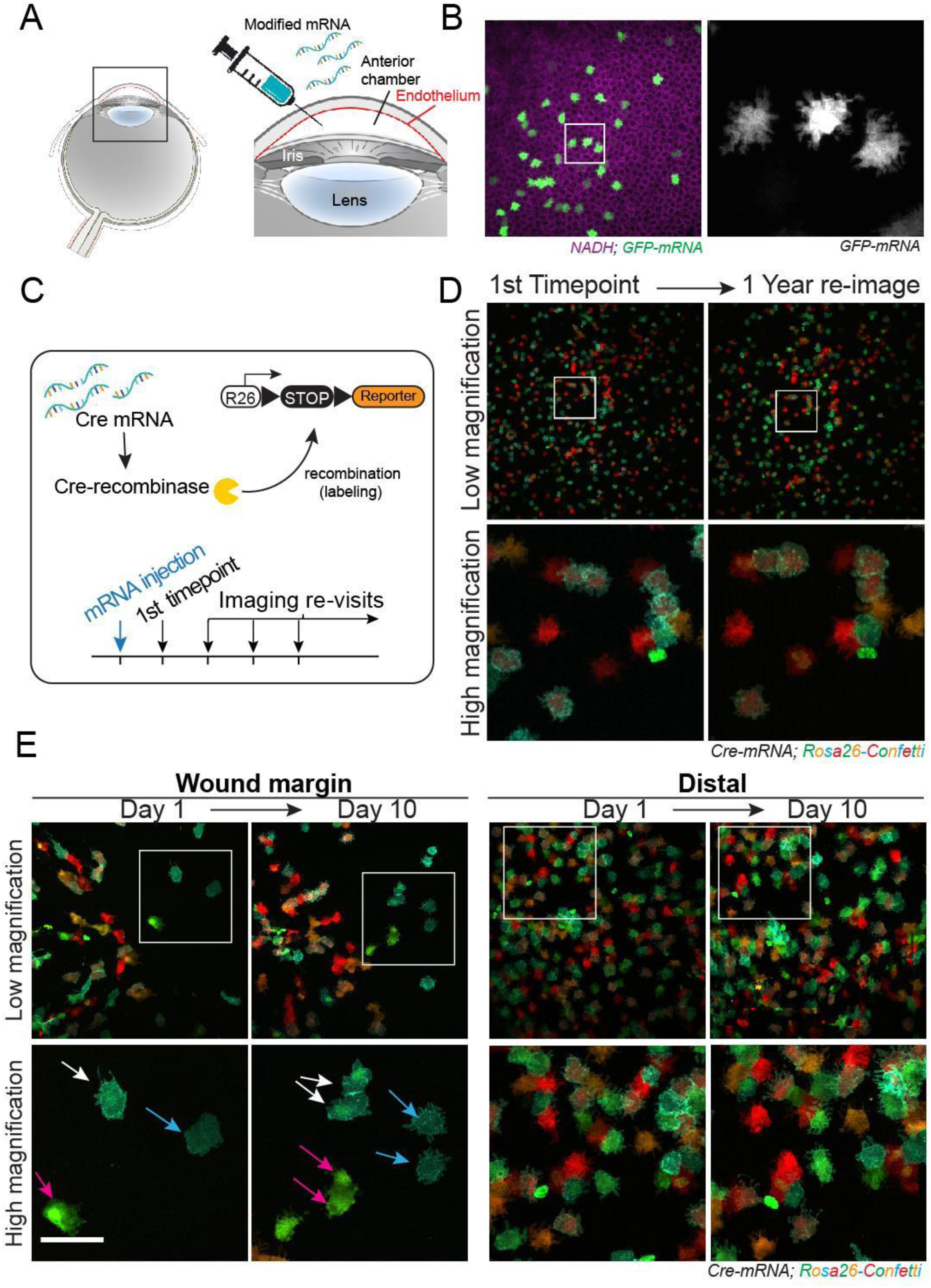
Corneal endothelial cells are equipotent in their proliferative capacity. A) Experimental design for exogenous protein expression in corneal endothelial cells *in vivo*, via intracameral injection of encapsulated modified mRNA. B) Example of GFP expression in corneal endothelial cells 24 hours after mRNA injection into the mouse eye. C) Experimental design of in vivo lineage tracing of corneal endothelial cells via in situ delivery of Cre-recombinase encoding mRNA, followed by longitudinal live imaging. D) Examples of multi-color *in vivo* lineage tracing of corneal endothelial cells, using the *Rosa26-confetti* reporter. The panel shows low and high magnification images of the same mouse eye imaged after mRNA injection (1^st^ Timepoint) and re-imaged after one year. E) Lineage tracing of corneal endothelial cell activity during injury repair. Left panels show low and high magnification views of the wound margins, taken at one and ten days after injury. Arrows show representative single corneal endothelial cells at the wound margin that divided during this timeframe. Right panels show equivalent areas of the same corneas located distally from the wound margin and illustrate the absence of cell divisions in these areas during wound healing. Scale bars 50 μm.

To track the activity of individual corneal endothelial cells under normal tissue physiology and to resolve their contribution to tissue repair, we performed *in vivo* lineage tracing by longitudinal live imaging. For this, we used mice harboring a fluorescent Cre reporter and induced recombination and clonal labeling by local injection of modified Cre-encoding mRNA (**Fig 3C**). Imaging of the corneas at 24 hours after injection confirmed Cre-induced recombination and labeling of corneal endothelial cells (**Fig 3D**). Furthermore, no new recombination events were observed after 24 hours, indicating that mRNA uptake is limited within this time window. Re-imaging the same corneas one year after labeling, we confirmed the near absolute quiescence of corneal endothelial cells, as the majority of labeled cells remained virtually unchanged in their original positions within the tissue (**Fig 3D**). Next, we tracked corneal endothelial cells during injury, to resolve their activities and contribution to the repair process. By imaging the corneas at daily intervals after the injury we captured profound changes in corneal endothelial cells located near the wound margins (**Fig S7**). These cells rapidly migrated towards the wound bed, becoming enlarged and extending filopodia-like structures (**Fig S6**). Through their concerted activity, the edges of the wound were brought together and adhered within three days after injury, effectively closing the wound. After this point, and once the continuity of the monolayer was re-established, no further changes were observed in the number, location, or morphology of individual cells within the corneal endothelium (**Fig S6**).

In our previous experiments, we demonstrated the existence of proliferating corneal endothelial cells during injury repair, using an *in vivo* cell cycle reporter (CyclinB1-GFP). However, due to the dynamic nature of this reporter we could only capture a snapshot of cells that are in the G2/M phase of the cell cycle at the time of imaging. To analyze the full extent of proliferation and resolve the history and spatiotemporal dynamics of mitotically active corneal endothelial cells, we developed a multi-color lineage tracing strategy to effectively track individual cells and their progeny with high accuracy (**Fig S7A**). After injection of the Cre-encoding mRNA, the corneas were imaged to document the location of labeled cells, and then re-imaged after two months, which confirmed that the tissue was fully quiescent, with no evidence of proliferation or other changes to the corneal endothelial cells during this time (**Fig S7B**). By tracking individual cells after induction of injury we present definitive evidence that corneal endothelial cells are activated and can undergo one or more cycles of cell division before returning into a quiescent state (**Fig 3E, S7B**). Importantly, we found that activation of corneal endothelial cells occurs exclusively at the wound margins, while the rest of the tissue is maintained in a quiescent state throughout the repair process and thereafter (**Fig 3E, S7B**). To investigate this further, we implemented an orthogonal approach, using *in vivo* photo-labeling as an unbiased way to track cells proximal and distal to the wound margin (**Fig S8**). In agreement with our previous data, only photo-labeled cells proximal to the wound responded to the tissue repair (**Fig S8**). These data suggest a ubiquitous regenerative potential of corneal endothelial cells *in vivo* that can be modulated by environmental signaling cues.

A promising therapeutic strategy to revert the disease progression and restore vision is to modulate intrinsic molecular pathways to promote corneal endothelial cell proliferation in the absence of any injury [29]. Modified mRNA technology has made it possible to transiently modulate the protein expression program of cells in vivo, offering an appealing vehicle for the implementation of this therapeutic strategy [30]. Our data of mRNA-mediated expression of exogenous proteins in corneal endothelial cells *in vivo*, empowered us to develop a therapeutic strategy to reprogram corneal endothelial cells into a pro-regenerative state without injury, and use our live imaging system to validate its efficacy. Our hypothesis is that by forcing exogenous expression of key enzymes and transcription factors we can activate the relevant signaling pathways to induce the corneal endothelial cells to undergo mitosis. To test this hypothesis, we performed a literature search and selected a combination of five pro-growth factors (<u>C</u>dk4, <u>C</u>cnd1, <u>M</u>yc, <u>S</u>ox2 & <u>Y</u>ap; henceforth abbreviated as CCMSY factors) (**Fig S9**). Cdk4 and Ccnd1(cyclin D1) typically form a complex that promotes progression of the cell cycle [29,30]. Myc and Sox2 - two of the Yamanaka factors for somatic cell re-programming - are known to promote proliferation in various cell types, including in human corneal endothelial cells *in vitro* and in a rat cryoinjury model *in vivo* [31,33]. Yap is a downstream effector of the Hippo pathway and critical for mediating contact inhibition growth, which is hypothesized to be a major driver of cellular quiescence in the corneal endothelium [34,35].

To test the efficacy of the CCMSY factors to induce proliferation of corneal endothelial cells *in vivo*, and to document any potential adverse effects, we performed concurrent lineage tracing of the cells that uptake and express the CCMSY mRNAs after eye injection (**Fig 4A**). To validate this assay, we first tested injecting a mix of different mRNAs that encode traceable reporters (GFP + mCherry or GFP + Cre-recombinase) and confirmed that cells which expressed one reporter always expressed the other. This indicates that a mix of encapsulated mRNA molecules is equally uptaken and expressed by the same corneal endothelial cells in vivo (**Fig S5C**). Thus, we used a Cre-encoding mRNA injected in a Cre reporter mouse line as a surrogate to label and track the corneal endothelial cells that express the CCMSY factors. At 24 hours post-injection in aged mice, cells that received the Cre + CCMSY mRNAs underwent mitosis (**Fig 4B-D, S10**). This spur of proliferation led to a measurable increase in corneal endothelial cell density effectively reversing the corneas of one-year-old mice to that of a three-month-old (**Fig 4E-H**). Control corneas that only received the Cre mRNA remained virtually unchanged with no evidence of proliferation, consistent with our previous lineage tracing experiments (**Fig 4B-E, S10**). Importantly, the pro-proliferative effect of the CCMSY treatment was only transient, with cells promptly returning to quiescence three days after mRNA injection (**Fig 4F**). Corneal endothelial cells that responded to the treatment with the CCMSY mRNAs maintained a normal morphology and expression of typical corneal endothelial markers, suggesting that their physiology was not altered, and the tissue remained stable two months after the injection (**Fig 4C, I; S10C**). We reproduced these results using a different mouse model (CyclinB1-GFP), which also demonstrated that activation of the cell cycle was specific to cells that expressed the CCMSY factors (**Fig 4D, S11**). It is important to note that in corneas which received the CCMSY mRNAs, proliferation was induced throughout the endothelium and in areas distal from the site of injection, suggesting that this was independent from potential injury cues. Moreover, no uncontrollable proliferation or other adverse effects were observed in subsequent re-imaging of the same corneas beyond the initial growth spurt (**Fig 4B-H, S10, S11**).

**Figure 4.**
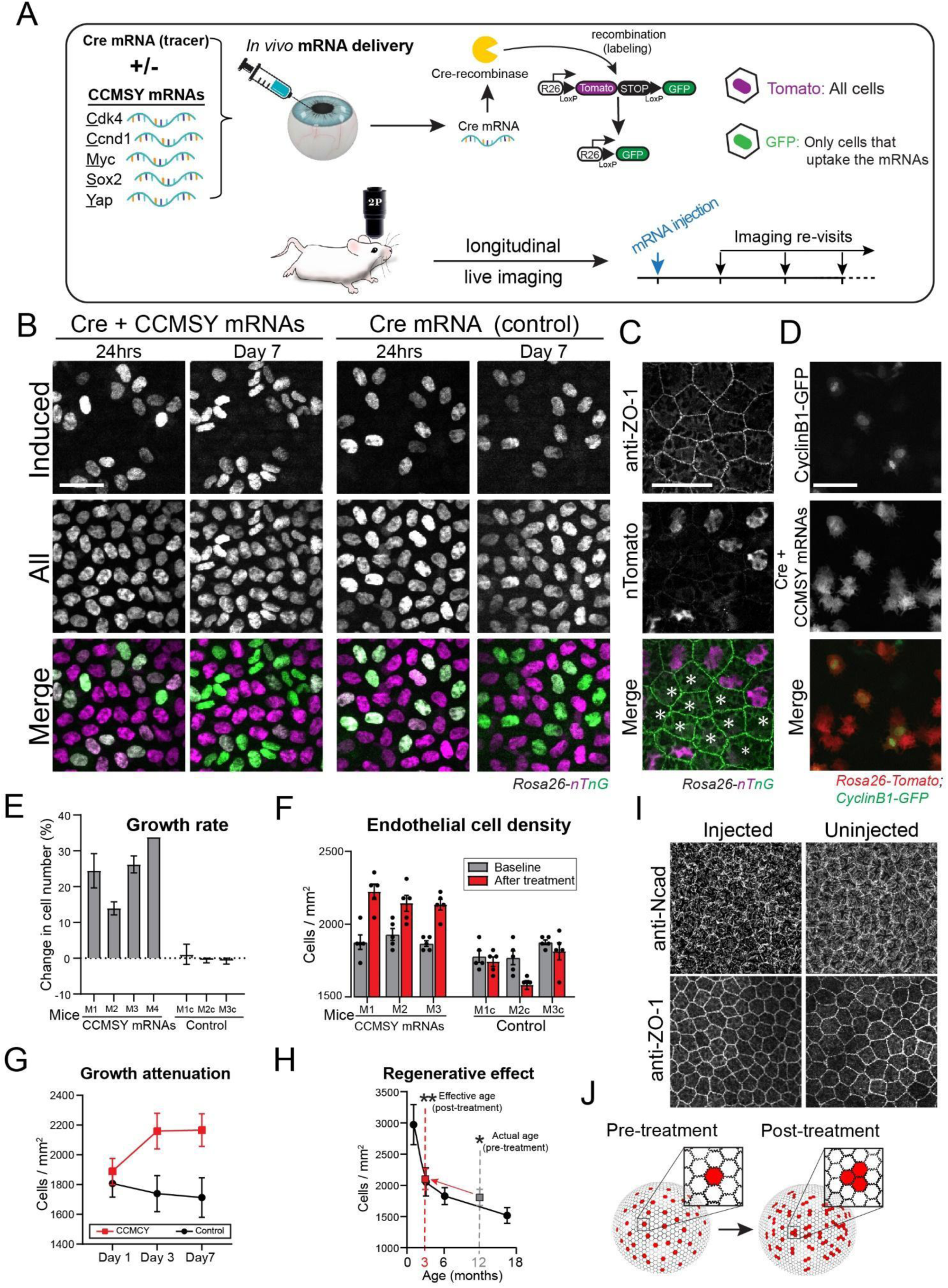
mRNA-mediated cell cycle activation of corneal endothelial cells *in vivo*. A) Experimental design for reprogramming and induced proliferation of corneal endothelial cells via intracameral injection of CCMSY mRNAs. A Cre-encoding mRNA is used to label cells that uptake the mRNAs in the corneas of a Cre-reporter mouse line (*Rosa26-nTnG*). Cells that incorporate the mRNAs switch from a nuclear Tomato to a nuclear GFP signal and their fate can be traced longitudinally. The activity of corneal endothelial cells is tracked over time by longitudinal live imaging. The same corneas were re-imaged at the indicated timepoints after mRNA injection. B) Representative examples from a lineage tracing experiment following mRNA injections. Cells that receive only the Cre mRNA remain quiescent while those that also receive the CCMSY mRNAs undergo cell divisions. C) Whole mount immunolabeling with the endothelial marker ZO-1, of corneas from *Rosa26-nTnG* mice collected ten days after injection with mRNA encoding for Cre plus the cocktail of pro-growth proteins. Asterisks show cells that have incorporated and expressed the mRNA as indicated but the switch from a nuclear-Tomato to a nuclear-GFP signal (*Rosa26-nTnG* reporter). D) Example of a lineage tracing experiment using the in vivo cell cycle reporter (*Cycling B1-GFP*) in combination with the Cre-reporter (*Rosa26-tdTomato*). Cells that receive the Cre + CCMSY mRNAs are the only ones that show positive GFP signal, indicating selective activation of the cell cycle in these cells. E) Graph showing the regenerative effect of the CCMSY mRNAs treatment. The average cell density of the corneal endothelium in 12-month-old mice injected with the CCMSY mRNAs increased after treatment to levels equivalent to those of 3-month-old mice. For quantification of cell density as a function of age see Figure 1C. F) Quantification of change in total endothelial cell density of corneas injected with either the Cre-recombinase alone (Control), or plus the CCMSY mRNAs. The same corneas were re-imaged at the indicated timepoints. Each bar pair on the graph represents a different cornea imaged before and after treatment [N = 5 measurements per cornea per timepoint; Cocktail: P = 0.0013(1), 0.0147(2), 0.0002(3); Control: P = 0.5 (1), 0.01(2), 0.33(3); paired t-test]. G) Quantification of change in the number of labeled cells that receive either the Cre-mRNA alone (Control), or Cre plus the CCMSY mRNAs. [N = 7 corneas, P < 0.0001; two-way ANOVA] H) Growth curves of the corneal endothelium in mice injected with the Cre-recombinase alone (Control), or plus the CCMSY mRNAs. Note that CCMSY induce only transient proliferation in the tissue. I) Whole mount immunolabeling with the endothelial markers zonula-occludens-1 (ZO-1) and N-cadherin, of corneas collected two months after injection with CCMSY mRNAs and compared non-injected. J) Schematic of the mRNA-mediated proliferation of corneal endothelial cells. Endothelial cells that incorporate the mRNAs after the intracameral injection are shown in red. After treatment, endothelial cells that express the proteins encoded by the CCMSY mRNAs undergo a limited round of proliferation which leads to a stable increase in endothelial cell density. Scale bars 200 μm.

These experimental data provide a proof-of-concept that quiescent corneal endothelial cells can be reprogrammed and induced to proliferate *in vivo*, by modulating key molecular pathways, using modified mRNA delivered directly into the eye (**Fig 4J**). Despite the potential oncogenic nature of the factors tested here, the safety profile of modified mRNA-based therapies for corneal endothelial dysfunction is advantageous, due to the lack of genomic integration and inherent instability and short half-life of mRNAs in the cytoplasm. This is in line with our data showing a transient uptick in cell proliferation during the first 24 hours after injection, and a quick return to quiescence. The stability and performance of therapeutic mRNAs can be improved, by 5’ and 3’ sequence optimizations, synthetic capping, and incorporation of modified nucleosides [36–38]. High efficiency and improved specificity for the *in vivo* delivery of the mRNA cargo can also be achieved with the use of optimally formulated lipid nanoparticles and incorporation of cell-specific antibodies, as recently demonstrated [39]. These modifications can further minimize potential off-target effects of this treatment that went undetected in this study. Furthermore, due to the relative low risk of the treatment, dosing of the delivered mRNAs can be incremental and adjusted to achieve a desired outcome that restores the corneal endothelium to a functionally optimal cell density. While many of these improvements and modifications were incorporated into this study, future work is needed to systematically test these parameters to achieve ideal clinical outcomes. We envision that the *in vivo* models and mRNA-based methodologies developed for the purposes of this and other recent studies will serve as a paradigm for basic research and enable new therapeutic platforms to address unmet clinical needs for corneal endothelial and other ocular diseases [40]. Proliferation-inducing strategies using modified mRNA technology may also have far-reaching applications for other organs affected by cellular attrition due to aging or disease.

## Acknowledgments

We thank George Cotsarelis, Ken Zaret, Jonathan Epstein and Jean Bennet for their feedback and critical review of this manuscript. This study was partially supported by a seed grant from Penn Health Tech and the Regenerative Ophthalmology Program, made possible by a generous donation from Richard and Carolyn Sloane, as well as NIH/NEI (R01EY016134, P30ET12576, and K08EY028199). The sequencing was carried by the DNA Technologies and Expression Analysis Core at the UC Davis Genome Center, supported by NIH Shared Instrumentation Grant 1S10OD010786-01. We also acknowledge the support of the Institute for Regenerative Medicine and the entire stem cell community at Penn. P.R. was supported by grants from NIH/NEI (R01EY030599) and from the American Cancer Society (RSG1803101DCC). S.T. was supported by a grant from NIH/NEI (R01EY016134). B.L. was supported by a grant from NIH/NEI (K08EY028199).

## Author contributions

P.R. conceptualized the study, designed and performed the experiments. M.M. performed the breeding and genotyping of mice. L.O. and B.S. performed histological analyses. T.P and H.P. assisted with the mRNAs IVT and Western blot analysis. R.R, S.P., B.C.L. and S.T. performed the single-cell RNA sequencing analyses. All authors discussed results and participated in the manuscript preparation and editing. P.R. supervised the study and wrote the manuscript.

## Competing interests

A provisional patent application has been filed for “Compositions and methods for regenerating corneal endothelium”.

## Data and materials availability

Data sets and reagents presented in this study are available from the corresponding author upon request.

## Figure Legends

**Supplemental Figure 1.**
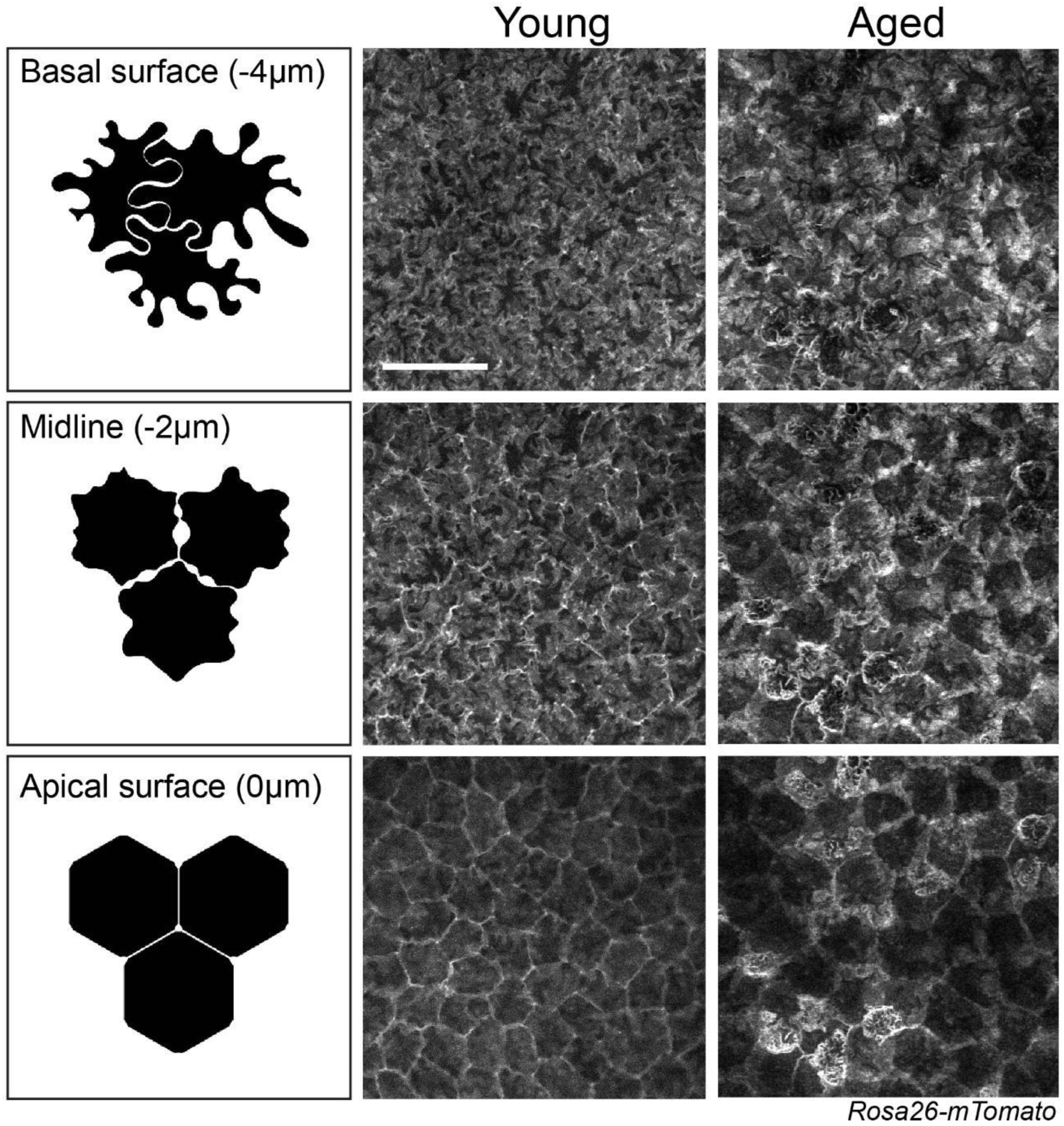
High resolution live imaging of the basal and apical surfaces of the corneal endothelium reveals differences in young and aged mice. Images were acquired using 2μm thick serial optical sections. Corresponding models illustrate the distinct cell-cell contacts and membrane organization of the basal and apical surfaces of the corneal endothelium (left panels; adapted from *He et al. Sci Rep 2016*). A globally expressed, membrane-localized fluorescent reporter (*Rosa26-mTomato*) was used to resolve the fine tissue morphology *in vivo*. Scale bar 50 μm.

**Supplemental Figure 2.**
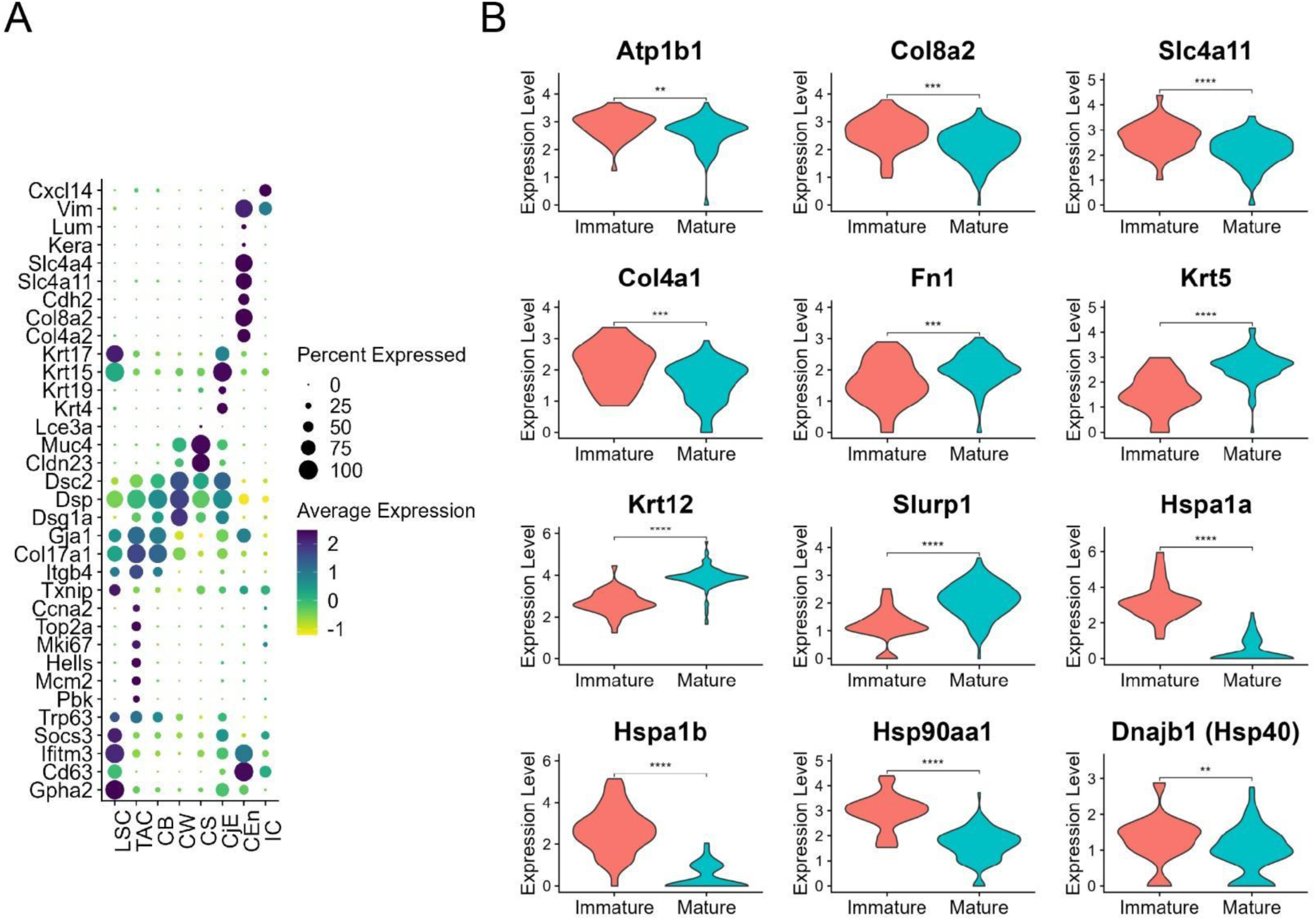
Aging corneal endothelial cells lose their identity. Unbiased clustering of integrated samples across three ages (2-, 6- and 11-month-old) in wild type mice revealed 8 cell populations of the cornea. (A) Dot plot demonstrating cell type specific marker expression patterns. (B) Violin plots of select differentially expressed genes of biological significance by maturity demonstrate decreased endothelial markers and increased epithelial, fibroblast, and senescence markers with age.

**Supplemental Figure 3.**
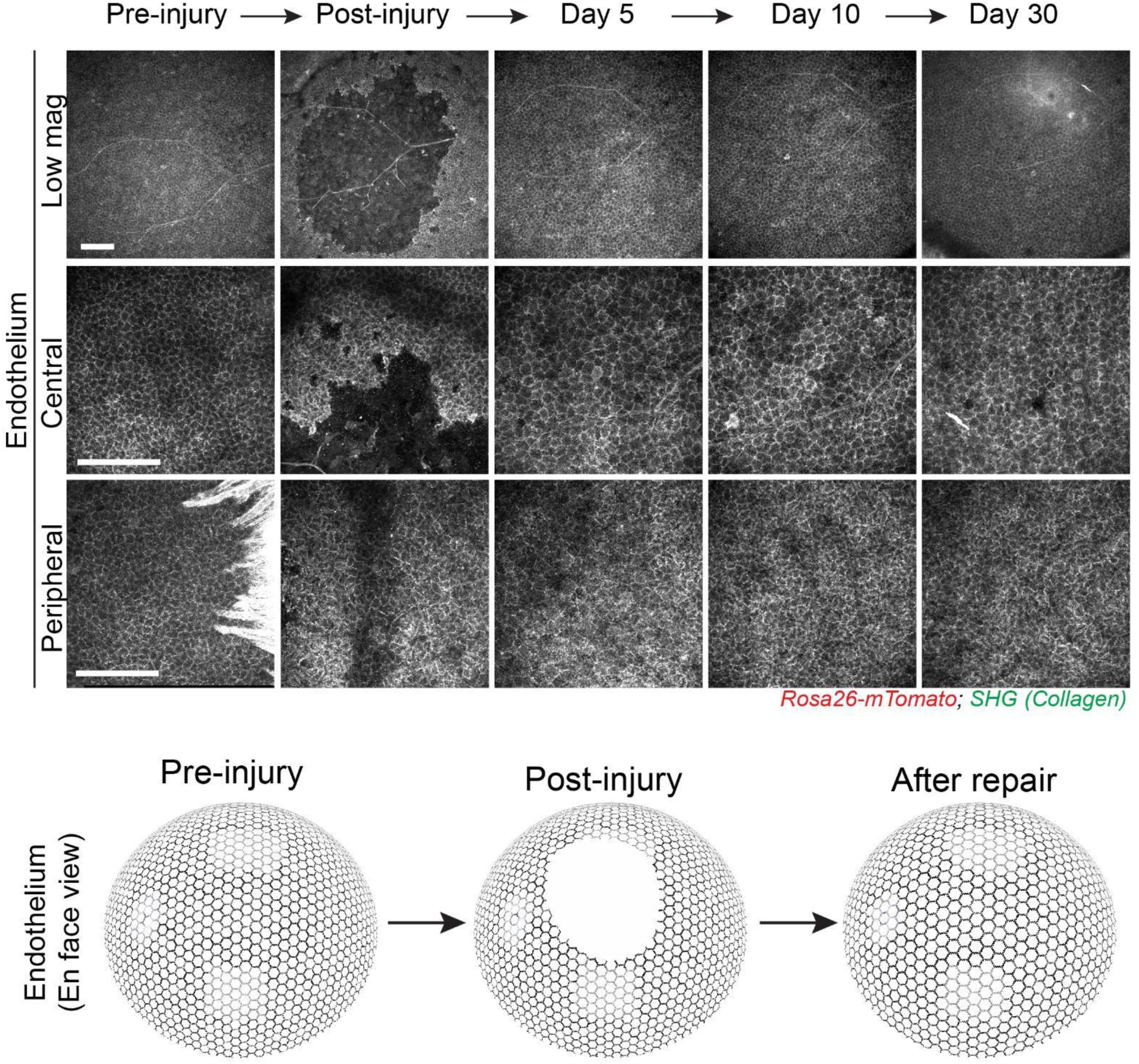
Wound healing re-establishes the corneal endothelial layer but results in reduced cell density. Representative low magnification maximal projections of the endothelial layers (en face view) and corresponding high magnification views of the central and peripheral corneal endothelium (bottom panels, en face view), taken at the indicated time points before and after injury. A globally expressed, membrane-localized red fluorescent reporter (*Rosa26-mTomato*) was used to resolve tissue and cell morphology *in vivo*. The stromal extracellular matrix is visualized by Second Harmonic Generation (SHG) and shown in green (top panel). Scale bars 200 μm.

**Supplemental Figure 4.**
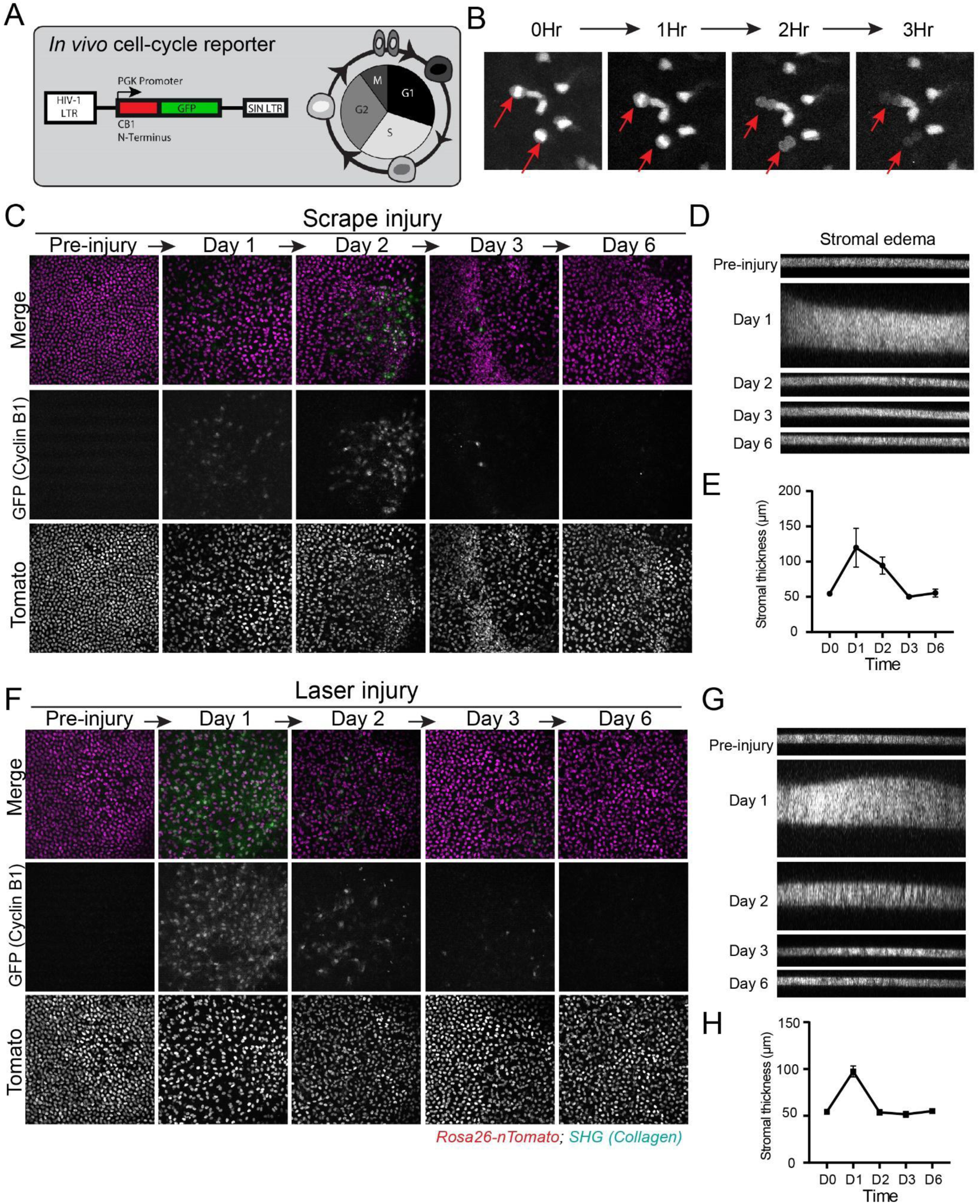
Longitudinal live imaging reveals the cellular and tissue dynamics of corneal endothelial injury response. A) Validation of the cell cycle reporter (*CyclinB1-GFP*) used to capture corneal endothelial proliferation in vivo. Top left panel shows the sequence elements of the transgenic reporter. Expression of GFP recapitulates the kinetics of the endogenous cyclin B1 enzyme, with a peak signal intensity during the G2/M phases of the cell cycle. B) Panels show representative frames from a time sequence, which depict actively cycling cells in the live corneal epithelium. Red arrows show cells that complete mitosis, in whose daughter cells the GFP signal quickly becomes undetectable. C) Representative low magnification en face views of the corneal endothelium taken at indicated timepoints during scrape injury repair. Proliferating corneal endothelial cells can be visualized by green fluorescence (*CyclinB1-GFP*). D) Corresponding side views of the entire thickness of the cornea taken at the indicated time points before and after scrape injury. The stromal extracellular matrix is visualized by Second Harmonic Generation (SHG; XZ views). E) Graph showing changes in stromal thickness during scrape injury repair. F) Representative low magnification en face views of the corneal endothelium taken at indicated timepoints during femtosecond laser injury repair. Proliferating corneal endothelial cells can be visualized by green fluorescence (*CyclinB1-GFP*). G) Corresponding side views of the entire thickness of the cornea taken at the indicated time points before and after laser injury. The stromal extracellular matrix is visualized by Second Harmonic Generation (SHG; XZ views). H) Graph showing changes in stromal thickness during laser injury repair. Scale bars 200 μm.

**Supplemental Figure 5.**
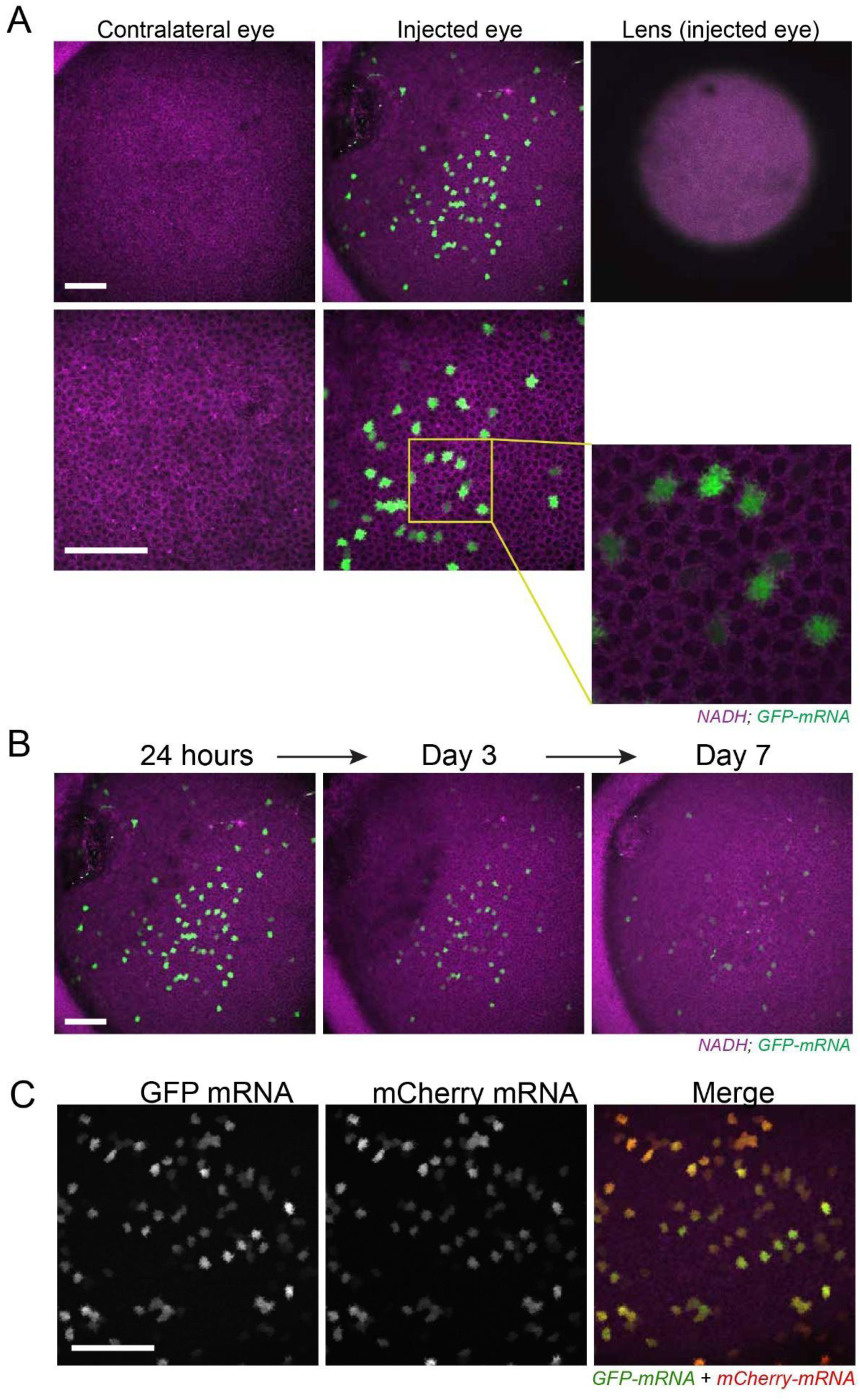
*In vivo* injection of encapsulated modified mRNA modulates protein expression in the murine corneal endothelium. A) Validation of efficacy and specificity of protein expression in the corneal endothelium by modified mRNA delivered *in vivo*, by injection into the anterior chamber of the eye (intracameral). A GFP encoding modified mRNA was injected into the left eye of a live mouse and the same eye was imaged 24 hours after injection. Expression of GFP was detected in the corneal endothelium of the injected. No GFP signal was detected in other cellular compartments of the cornea, the lens or anywhere in the un-injected contralateral eye. Two-photon imaging using 750 nm laser wavelength was used to generate metabolite autofluorescence (NADH) and was used as a means to visualize unlabeled cells in magenta. B) Representative low magnification en face views of the same corneas taken at the indicated timepoints. Expression of GFP wanes after 24 hours and detectable signal persists up to seven days from injection of the mRNA. C) Co-expression of GFP and mCherry in corneal endothelial cells after injection of a cocktail consisting of equal amounts of mRNA encoding for each fluorescent protein. Scale bar 50 μm.

**Supplemental Figure 6.**
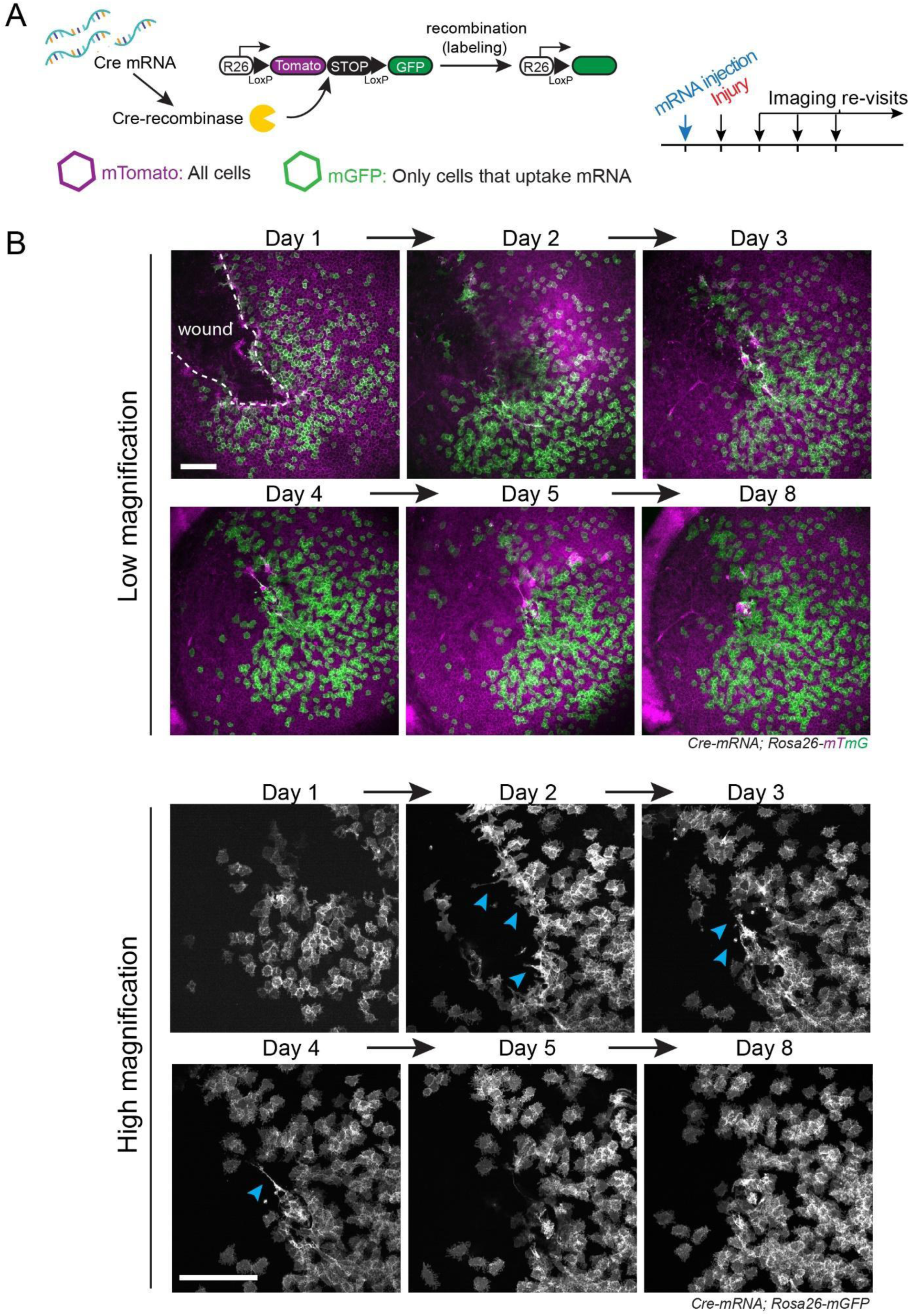
Cell migration contributes to wound healing of the corneal endothelium. A) Experimental design to visualize the activity of single endothelial cells and analyze their contribution to injury repair by in vivo lineage tracing. Clonal labeling of single corneal endothelial cells is achieved with a single injection of modified mRNA encoding for Cre-recombinase. Cells that express the enzyme are permanently labeled with the Cre reporter (*mGFP*) and their activity is tracked by longitudinal live imaging. B) Representative low (top panels) and high (bottom panels) magnification views of the wounded corneal endothelium taken at the indicated time points during injury repair. Cre-recombinase encoding mRNA was injected before the injury to mark corneal endothelial cells and track their activity during repair. A globally expressed, membrane-localized cre reporter (*Rosa26-mTmG*) was used to resolve tissue and cell morphology in vivo and to mark cells which express the Cre-recombinase with mGFP. Blue arrowheads indicate filopodia-like structures of migrating corneal endothelial cells during wound healing. Scale bars 200 μm

**Supplemental Figure 7.**
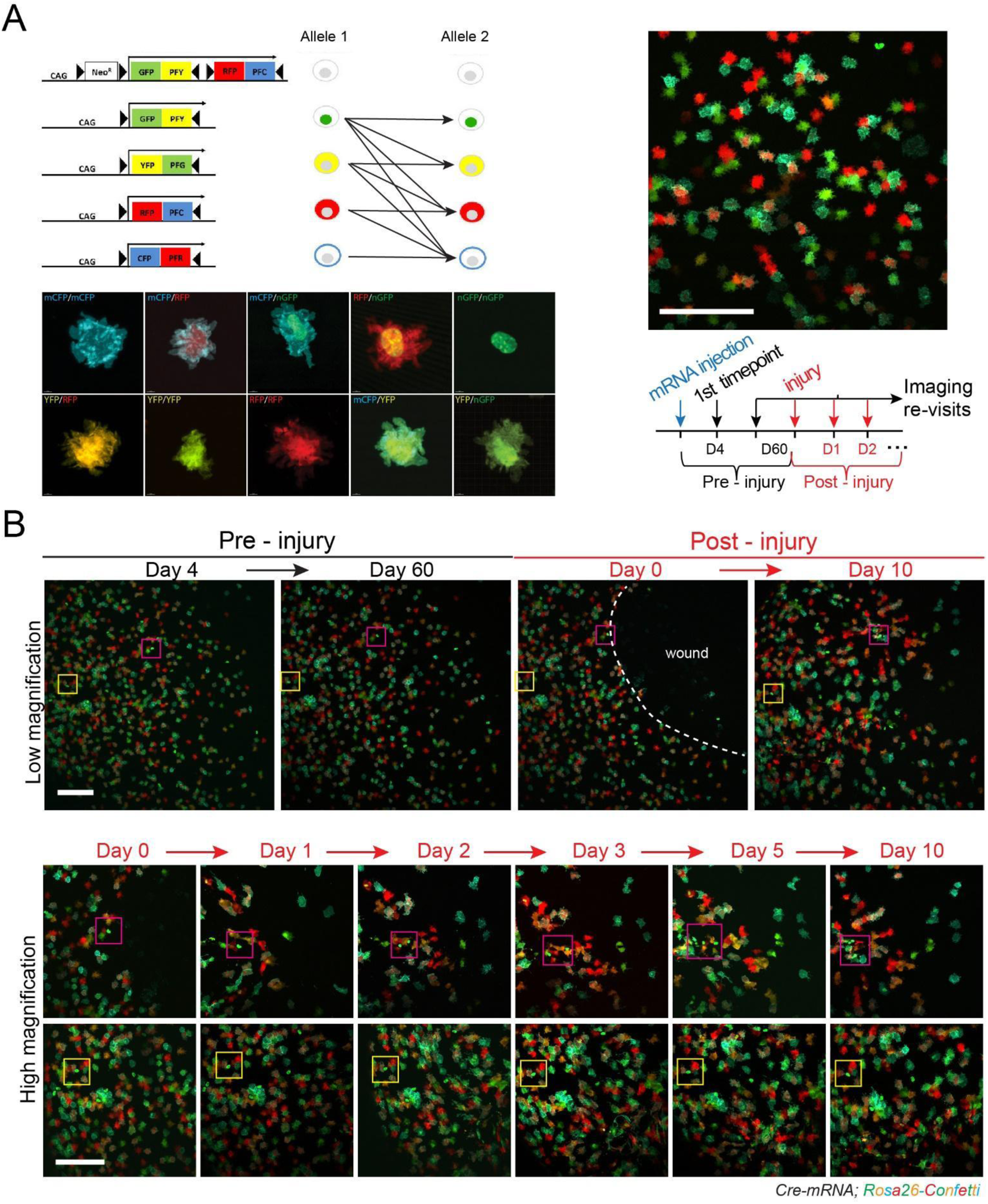
Multicolor *in vivo* lineage tracing reveals the distinct activities of corneal endothelial cells during wound healing. A) Validation of multi-color labeling of corneal endothelial cells. Diagram shows the sequence elements of the *Rosa26-confetti* reporter and the possible recombination events that result in ten unique color combinations when mice homozygous for the two confetti alleles are used. Imaging of the corneal endothelium after injection of Cre-encoding modified mRNA confirmed these theoretical color labels expressed stochastically in single corneal endothelial cells. B) Representative low (top panels) and high (bottom panels) magnification views of the corneal endothelium taken at the indicated time points before and after injury. Cre-recombinase encoding mRNA was injected before the injury to mark corneal endothelial cells and track their activity during repair. Note that labeled cells remain quiescent for at least two months before injury (yellow squares), but only those proximal to the wound margin become activated and proliferate after injury (red squares). Scale bars 200 μm.

**Supplemental Figure 8.**
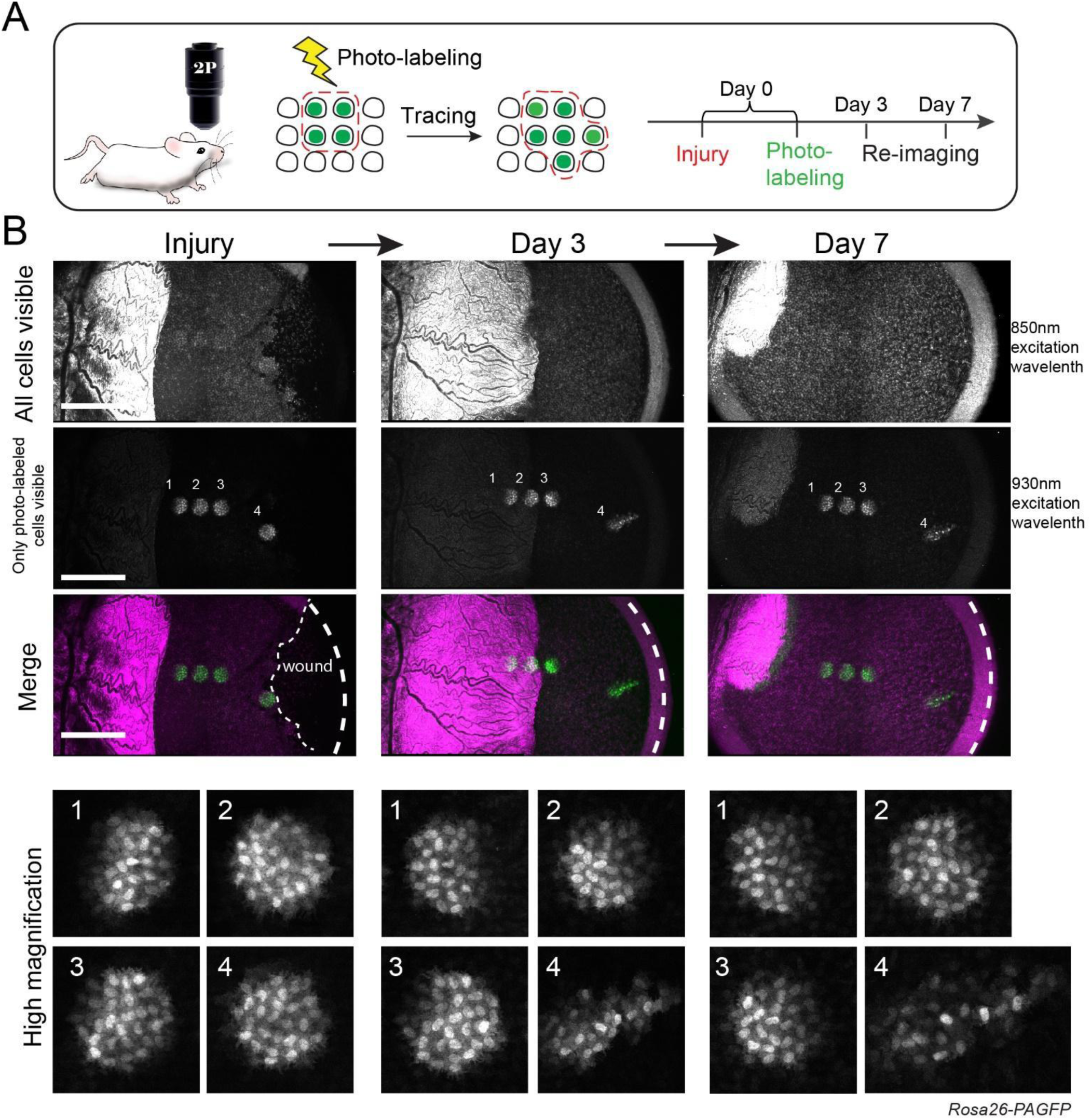
Unbiased *in vivo* lineage tracing captures exclusive contribution of corneal endothelial cells at the wound edge. A) Experimental strategy for un-biased lineage tracing of corneal endothelial cells by in vivo photo-labeling and longitudinal live imaging. B) Representative low (top panels) and high (bottom panels) magnification views of the corneal endothelium taken at the indicated time points after injury. Equivalent groups of cells were photo-labeled at various distances from the center of the cornea (1-3) as well as proximal to the wound margin (4). Imaging at 850 nm was used to visualize the pre-activated state of the globally expressed PAGFP reporter and was used as a counterstain to visualize unlabeled cells in the tissue. Imaging at 930 nm illuminates only cells in which the PAGFP was irreversibly photo-activated. The same corneas were re-imaged to track changes in the labeled cells after injury. Note that labeled cells distal to wound edge remain quiescent and do not directly contribute to injury repair. Scale bars 500 μm.

**Supplemental Figure 9.**
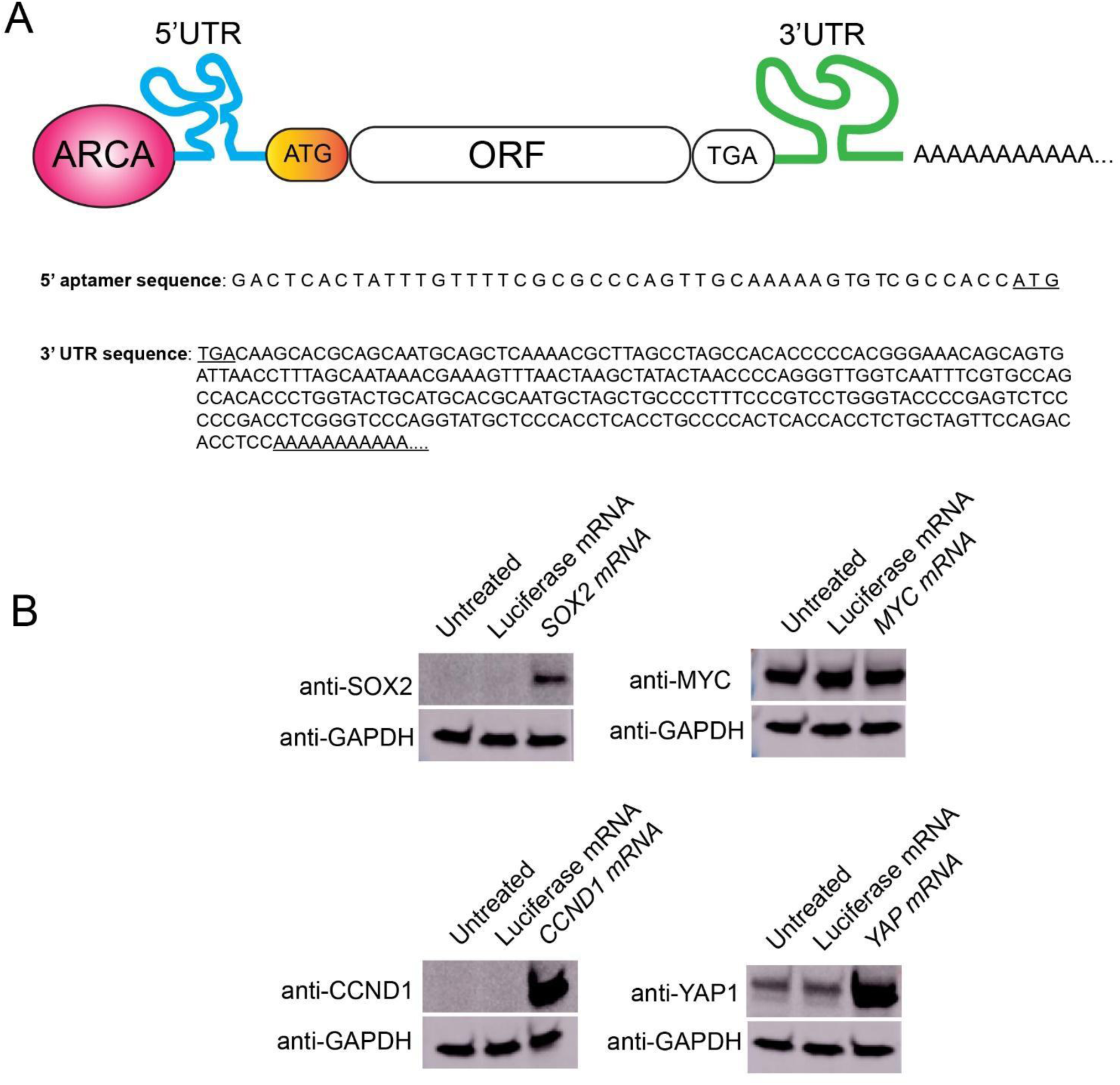
Validation of modified mRNA constructs. A) Sequence elements of mRNAs used in this study. See Material and Methods for more information. B) Western blot analysis of protein expression in 392T cells transfected with the indicated mRNAs.

**Supplemental Figure 10.**
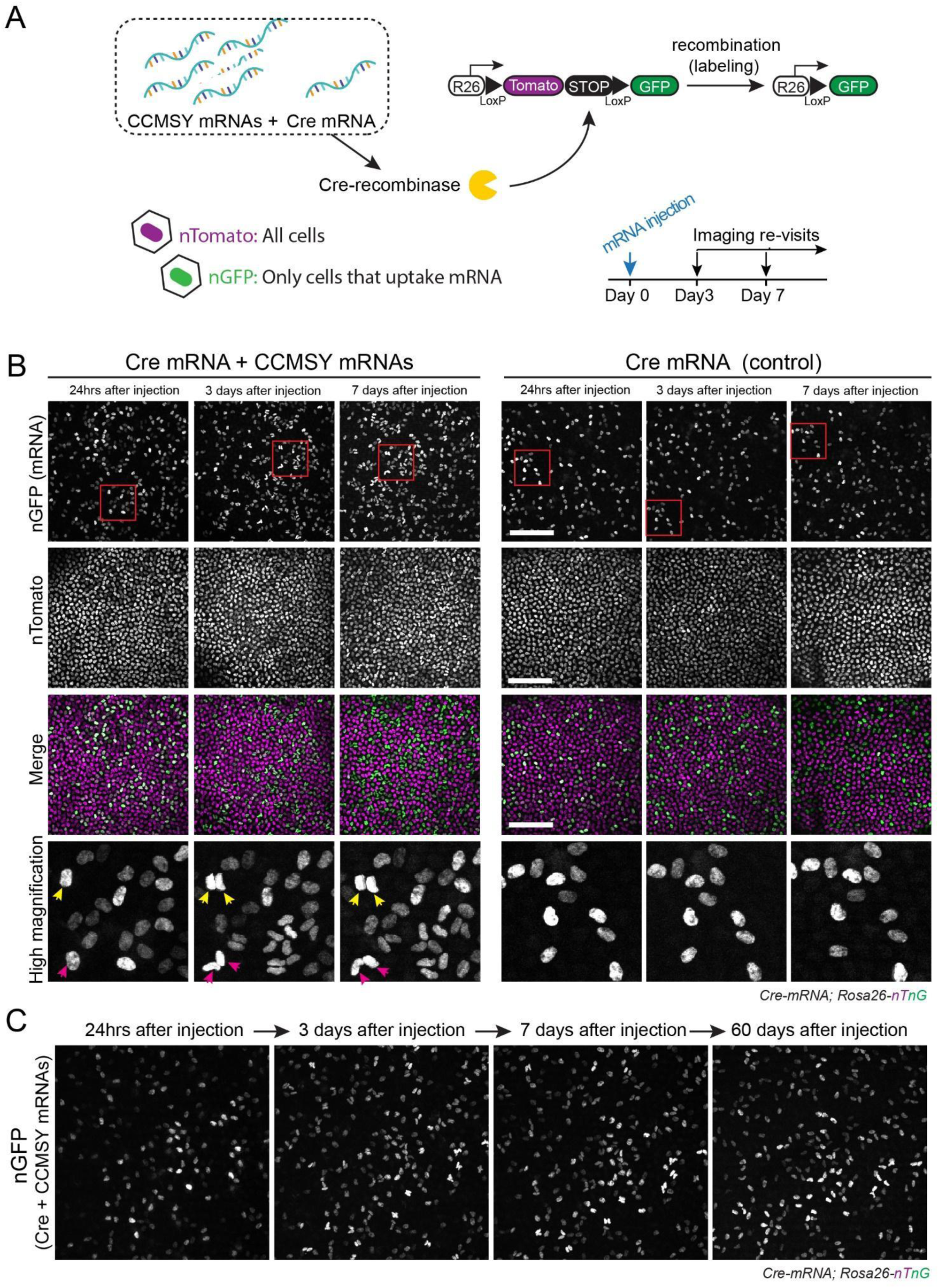
*In vivo* lineage tracing demonstrates efficacy of mRNAs delivered *in situ,* to elicit transient cell proliferation in the corneal endothelium. A) Experimental design for reprogramming and induced proliferation of corneal endothelial cells via in vivo injection of mRNAs encoding for pro-growth factors (<u>CCMSY</u>: <u>C</u>dk4 / <u>C</u>cnd1 / <u>M</u>yc / <u>S</u>ox2 / <u>Y</u>ap). A Cre-encoding mRNA is used to label cells that uptake the mRNAs in the corneas of a Cre-reporter mouse line (*Rosa26-nTnG*). The activity of corneal endothelial cells is tracked over time by longitudinal live imaging. B) Panels show representative low and high magnification en face views of the corneal endothelium at the indicated timepoints from mRNA injection of Cre-recombinase alone (Control), or plus the CCMSY mRNAs encoding for pro-growth proteins. Cells that incorporate the Cre-mRNA switch from a nuclear Tomato to a nuclear GFP signal (*Rosa26-nTnG*) and their fate is traced longitudinally by live imaging. Red squares indicate the same cells imaged during the time course and depicted in high magnification in the bottom panels. Yellow and red arrowheads indicate examples of cells that are marked with nuclear-GFP after the uptake and expression of Cre mRNA (+CCMSY mRNAs), undergoing a cell division three days after injection and returning to quiescence thereafter. C) Long-term follow-up represented in the time-series of the corneal endothelium taken at the indicated timepoints from mRNA injection with CCMSY+Cre mRNAs. Note that after the initial spur of proliferation the tissue remains stable for at least two months after the time of injection. Scale bars 200 μm.

**Supplemental Figure 11.**
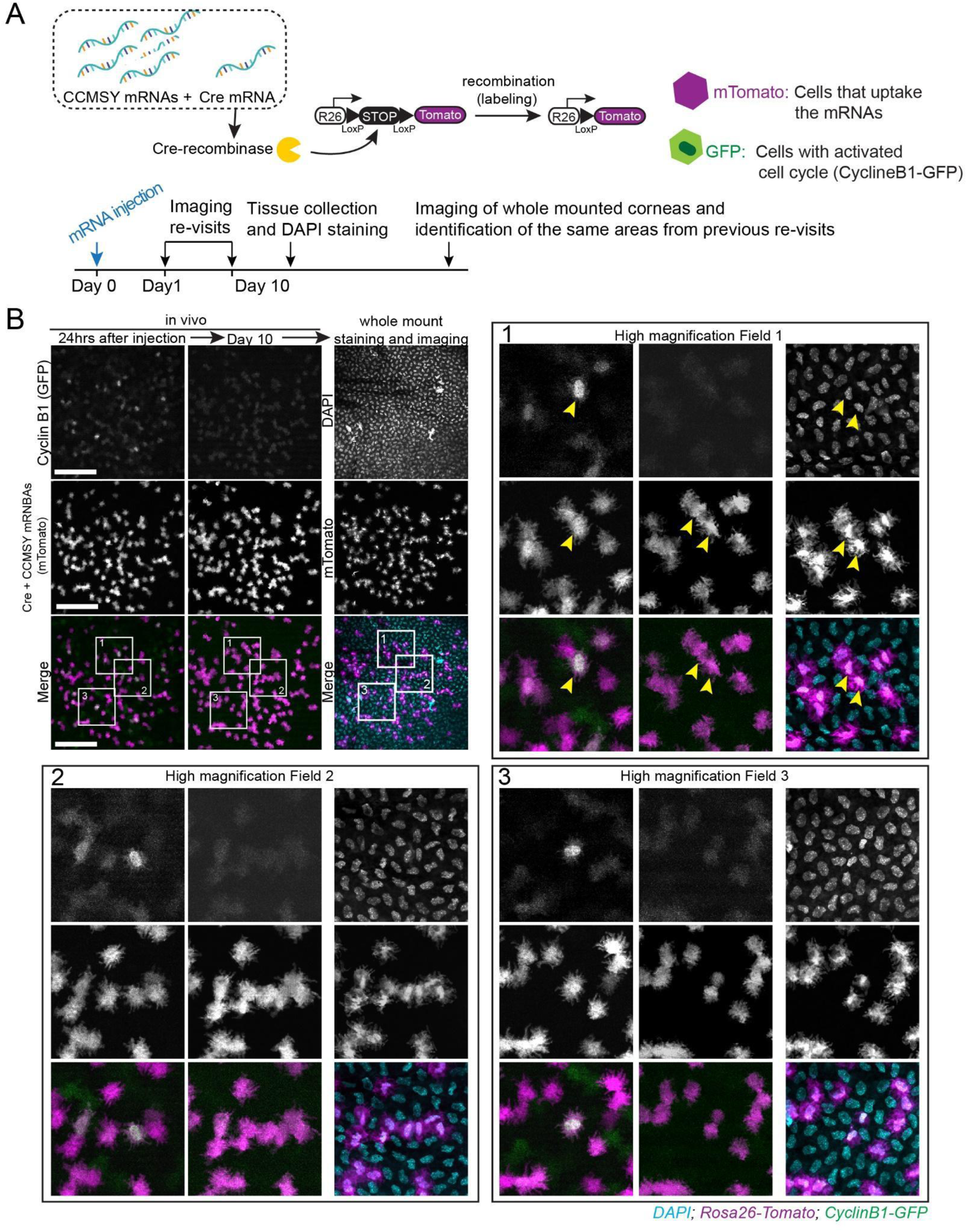
*In vivo* cell cycle reporter is specifically activated in corneal endothelial cells that uptake the mRNAs. A) Experimental design for reprogramming and induced proliferation of corneal endothelial cells via in vivo injection of mRNAs encoding for pro-growth factors (<u>CCMSY</u>: <u>C</u>dk4 / <u>C</u>cnd1 / <u>M</u>yc / <u>S</u>ox2 / <u>Y</u>ap). The mice used in these experiments express an in vivo cell cycle reporter (*CyclingB1-GFP*). A Cre-encoding mRNA is used to label cells that uptake the mRNAs in the corneas of a Cre-reporter mouse line (*Rosa26-nTomato*). Lineage tracing was performed to analyze induced proliferation of corneal endothelial cells in vivo, following intracameral eye injection of CCMSY mRNAs. The same corneas where imaged 24 hours after mRNA injection and again ten days later. After the final imaging the corneas where dissected, stained with DAPI to visualize the nuclei of corneal endothelial cells and imaged again as whole mounts. B) Panels show representative low and high magnification en face views of the corneal endothelium at the indicated timepoints from CCMSY+Cre mRNAs injection. White squares indicate the same cells imaged during the time course and depicted in high magnification in panels 1-3. Yellow arrowheads indicate one example of a cell that is marked with tdTomato after the uptake and expression of Cre mRNA (+CCMSY mRNAs), also showing positive Cyclin B1-GFP signal. Scale bars 200 μm.

## Methods

### Mice

All procedures involving animal subjects were performed with the approval of the Institutional Animal Care and Use Committee (IACUC) of the University of Pennsylvania or University of California, Davis and were consistent with the guidelines set forth by the ARVO Statement for the Use of Animals in Ophthalmic and Vision Research. *R26^loxp-stop-loxp-tdTom^*, *R26^loxp-tdTom-stop-loxp-EGFP^*, *R26^loxp-nTom-stop-loxp-nGFP^*, *R26^lox-stop-loxp-Confetti^* and *Pgk1CyclinB1^-GFP^* mice were obtained from The Jackson Laboratory. *R26^PAGFP^* reporter mice were generated by crossing the *E2a^Cre^* with *R26^loxp-stop-loxp-PAGFP^*lines, obtained from The Jackson Laboratory, to achieve germline transmission of the recombined allele and ubiquitous expression of the PAGFP reporter. All mice that were used in this study were bred for multiple generations into a Crl:CD1(ICR) mixed background. Mice were housed in a temperature and light-controlled environment and received food and water *ad libitum*. Up to 5 mice of the same sex and similar age were housed in a cage. Mice were provided Bed-o’Cobs (The Andersons Lab Bedding), a porous cob material, as bedding and Shred-n’Rich nestlets (The Andersons Lab Bedding) for nesting and enrichment.

### Intravital imaging of the corneal endothelium

Preparation of the mice for intravital imaging of the eye was performed with the following amendments to the previously described protocol (Rompolas et al. 2016). Mice were initially anesthetized with IP injection of ketamine/xylazine cocktail (0.1 ml / 20 g body weight; 87.5 mg / kg Ketamine, 12.5 mg / kg Xylazine). A deep plane of anesthesia was verified by checking pedal reflexes. The mouse head was stabilized with a custom-made stereotaxic apparatus that includes palate bar and nose clamp but no ear bars. Precision, 3-axis micro-manipulators are used to adjust the head tilt so that the eye to be imaged is facing up. A drop of eye gel (0.3 % Hypromellose) was used as an optically neutral interface between the eye and a glass coverslip, and to prevent dryness and irritation to the tissue during the anesthesia and imaging procedure. After preparation and mounting is complete, the stage is placed on the microscope platform under the objective lens. A heating pad is used to keep a stable body temperature and vaporized isoflurane is delivered through a nose cone to maintain anesthesia for the duration of the imaging process. After each imaging session, the eyes were rinsed with PBS and the mice were monitored and allowed to recover in a warm chamber before returned to the housing facility.

### Imaging equipment and acquisition settings

Image acquisition was performed with an upright Olympus FV1200MPE microscope, equipped with a Chameleon Vision II Ti:Sapphire laser. The laser beam was focused through 10X, 20X or 25X objective lenses (Olympus UPLSAPO10X2, N.A. 0.40; UPLSAPO20X, N.A. 0.75; XLPLN25XWMP2, N.A. 1.05) an emitted fluorescence was collected by two multi-alkali and two gallium arsenide phosphide (GaAsP) non-descanned detectors (NDD). The following wavelengths were collected by each detector: NDD1 419-458 nm; NDD1 458–495 nm; GaAsP-NDD1 495–540 nm; GaAsP-NDD2 575–630 nm. GFP and Tomato reporters were excited at 930 nm and their signal was collected by GaAsP-NDD1 and GaAsP-NDD2, respectively. Second harmonic generation (SHG) signal was generated using 850 nm or 930 nm excitation wavelengths and detected by NDD1 or NDD2, respectively. Serial optical sections were acquired in 2–5 μm steps, starting from the surface of the eye and capturing the entire thickness of the cornea (epithelium ∼40 μm, stroma/endothelium ∼80 μm). Expanded views of the cornea and limbus were obtained by acquiring a grid of sequential optical fields-of-view that were automatically stitched into one high-resolution tiled image using the microscope manufacturer software. Multi-day tracing experiments were done by re-imaging the same field-of-view or the entire eye at the indicated times after the initial acquisition. For each time point, inherent landmarks within the cornea, including the organization of the vasculature and collagen fibers (SHG), were used to consistently identify the limbus and navigate back to the original regions. Macroscopic images of the mouse eye were acquired under brightfield and fluorescence with an Olympus MVX10 Fluorescent Macro Zoom microscope fitted with Hamamatsu Orca CCD camera for digital imaging.

### Photo-labeling

Photo-labeling experiments with the *R26^PAGFP^* reporter mice were carried out with the same equipment and imaging setup as used for acquisition. The pre-activated form of the fluorescent proteins was visualized by exciting with 850 nm wavelength and emission signal was collected in GaAsP-NDD1 (495-540 nm). Excitation with 930 nm verified that no signal is emitted by the reporters before activation. Photo-labeling was achieved by scanning a defined region-of-interest (ROI) at the plane of the basal layer of the epithelium, with the laser tuned to 750 nm wavelength, for 5-10 sec, using 5-10% laser power. The z-plane was then moved down to the corneal endothelium, which served as a reference, and the same ROI was used for photo-labeling cells in that layer. Immediately after photo-activation, a series of optical sections, with a range that includes the entire thickness of the cornea, were acquired using the same acquisition settings as for GFP. Visualizing the signal of the activated form of PAGFP only within the ROI confirmed the successful photo-labeling of basal epithelial or endothelial cells. Following the initial image acquisition immediately after photo-labeling the same eyes were re-imaged at the indicated times to evaluate the changes of the labeled epithelial population and their movements compared to the endothelial reference cell group.

### Wounding assay

Corneal epithelial debridement wounds were generated as previously described (Chan & Werb 2015). Mice were initially anesthetized with IP injection of ketamine/xylazine cocktail and their eyes imaged under brightfield and fluorescence microscopy. Mice where then placed on a heating pad and observed under an Olympus SZ61 dissecting microscope. After topical application of Proparacaine the epithelium from the central part of the cornea was removed with an Algerbrush II ophthalmic brush to generate a scrape wound of 1.5 mm in diameter. The eyes were imaged again before the mice were allowed to recover.

### Intracameral eye injections

Adult mice ranging from 3-22 months of age mice were used for this procedure. For analgesia 1-2 drops of topical .5% proparacaine eye drops were instilled to each eye before the procedure. Mice were initially anesthetized with IP injection of ketamine/xylazine cocktail and kept warm using a temperature-controlled heating pad throughout the whole procedure. A deep plane of anesthesia was verified by checking pedal reflexes. The mouse head was stabilized by means of a palate bar and nose clamp and using a special sterotaxic apparatus that does not require the use of ear bars. Using precision micro-manipulators, the head angle was adjusted so that the treated eye was facing upwards. The pupils were dilated with .5% tropicamide ophthalmic drops and the treated eye was examined under a stereomicroscope to ensure that the pupil is fully dilated and that the ocular muscles are relaxed so that there is no eye movement. The absence of eye movements ensures stability during the injection. A diluted betadine solution was applied topically using a 1ml syringe to the ocular surface and fornices. For the injection a custom-made microinjection apparatus was used that attaches directly to the same heated mounting platform that is used for intravital imaging. The microinjector is composed of a hydraulic pump (Eppendorf CellTram Oil) connected to a 1mm inner diameter PTFE flexible tubing. A fitting adaptor connects the tubing to a single-use, sterile 31-gauge insulin pen needle (CareTouch). When attached to the tubing the microneedle is connected to a 3-axis manual micromanipulator to precisely control its movement during the injection. Using the microinjector pump a 5μl solution is loaded into the syringe, ensuring that no air bubbles enter the system or are present at the tip of the needle. Using the micromanipulators, the needle is guided to the mouse eye at a 45° angle, anteriorly relative to the limbus. The cornea is gently punctured so that the tip of the microneedle enters the anterior chamber, ensuring that the loaded microneedle remains at a 45° angle relative to the limbus during the puncture. Contact with the iris or the lens should be avoided. During this process the eye may need to be supported with a pair of disposable plastic tweezers from the opposite side. Using the microsyringe pump, ∼ 1.5 μl of solution is injected into the anterior chamber. This amount is injected gradually over 30 seconds before withdrawing the microneedle. Antibiotic ointment (Bacitracin Zinc and Polymyxin B Sulfate Ophthalmic Ointment USP) was applied to the treated eyes as prophylactic.

### Modified mRNA design, synthesis, and encapsulation

The constructs for the in vitro transcription (IVT) of cdk4, ccnd1, myc, sox2 and yap-encoding mRNAs were custom designed and synthesized by VectorBuilder using a template plasmid carrying a T7 promoter. A 5’ aptamer sequence for eIF4G binding was inserted between the T7 promoter and the start codon to optimize ribosome binding and translation initiation [38]. Downstream from the ORF and stop codon a 3’UTR sequence was inserted that consists of a combined 3’UTR from mitochondrial encoded 12S rRNA (mtRNR1) and the amino-terminal enhancer of split mRNA (AES), which has been reported to significantly improve mRNA stability and expression [41]. A minimum of 30 nucleotide long poly-A tail was also encoded into the DNA template. Template pDNA was linearized and ARCA-capped IVT-mRNA was produced using the T7 HiScribe kit from NEB, pseudo-uridine was incorporated into the reaction, and the product was Poly(a) tailed enzymatically using the manufacture’s recommendations. IVT-mRNA was then cellulose-purified to remove dsRNA products from the reaction. For validation of the IVT-mRNAs, 293T cells were cultured in 10% HI-FBS/1% PennStrep, then transfected with either luciferase IVT-mRNA or test IVT-mRNA. Cells were collected in cold DPBS and pelleted. Cell pellets were lysed with 1x RIPA + protease inhibitor cocktail. 40μg protein per sample was denatured in LDS reducing buffer containing BME at 70 degrees Celsius for 10 minutes. Protein was loaded into 4-12% Bis-Tris SDS Bolt gel and dry-transferred using iBlot instrument to nitrocellulose membrane. Blots were blocked in 5% milk/TBST overnight before incubating with primary and secondary anti-rabbit-HRP antibodies. Blots were developed using Immobilon ECL Ultra Western HRP Substrate and captured on Amersham Imager 600. Blots were stripped, blocked overnight, and re-probed with anti-GAPDH antibody. For control experiments, 5-methoxyuridine, nucleoside-modified CleanCap mRNAs encoding for EGFP, mCherry and Cre-recombinase were used (Trilink Biotechnologies). Encapsulation of modified mRNAs for intracameral injection was performed using Lipofectamine Messenger Max transfection reagent (Thermo Fischer) for following the manufacturer recommendations. A final concentration of 20 ng/μl of encapsulated mRNA in Opti-MEM medium (Thermo Fischer) was used for the in vivo injections.

### Preparation of single cell suspensions from 129S6/SvEvTac mice

Six corneas from 2 female and 1 male 129S6/SvEvTac mice of each age group (2, 6, and 11-month-old) were dissected and pooled in 600 µL of papain solution (CAT LK003178; Worthington Biochemical Corp., Lakewood, NJ) and incubated for 1.75 hours at 37 °C. The tube was inverted every 10-20 minutes during incubation to improve cell dissociation. The suspension was aspirated and passed through a Flowmi cell strainer (40 µM; CAT 136800040, Bel-Art, Wayne, NJ) and the strainer was rinsed with 300 µL of 1X PBS to collect additional adherent cells. The filtered suspension was centrifuged at 400 g for 10 minutes at 4 °C. The supernatant was discarded. The cell pellet was washed twice with 0.04% bovine serum albumin (BSA) in 1X PBS, centrifuging at 300 g for 5 minutes at 4 °C and the final pellet was resuspended on 40 µL of 2% BSA-PBS. Single cell suspensions for each age group were prepared in the same manner. All single cell suspensions were immediately submitted to the UC Davis DNA Technologies and Expression Analysis Core for 10X Chromium Single Cell 3’ library preparation and NovaSeq 4000 sequencing.

### Single cell RNA sequencing and analysis

The scRNA-sequencing was performed according to manufacturer’s recommendations. Raw FASTQ sequencing data was processed by the UC Davis Bioinformatics Core using 10X CellRanger for alignment to the 10x Genomics mouse reference genome. Analysis was then continued using Seurat v4 [44]. Count matrices for each age were converted to independent Seurat objects (2, 6 and 11MO) and quality control was performed through exclusion of cells with high or low gene counts, or high mitochondrial read percentage. Multiplet/doublet cells were excluded utilizing the package DoubletFinder [45]. Objects were scaled, normalized, and the top 2500 variable genes were identified for principal component analysis (PCA). Dimensionality of selection for Uniform Manifold Approximation and Projection (UMAP) was performed using Jack Straw and Elbow Plots. The three age objects were integrated according to the Seurat vignette [46]. Optimal clustering of cells was performed using the assistance of the clustree R package [47]. Cell type assignment was performed using marker genes defined in murine single cell literature [48–51]. Volcano plots were created using the R package EnhancedVolcano using differentially expressed gene lists generated by Seurat’s FindMarker function (Wilcoxon Rank Sum tests with P < 0.05) [52].

### Quantitative image analysis

Raw digital files from 2-photon imaging were acquired and saved in the OIB format using the microscope manufacturer’s software (FLUOVIEW, Olympus USA). To capture extended fields-of-view that encompass the entire ocular surface epithelium, including the cornea, limbus and conjunctiva, a tiling method was used to reconstruct a single image from multiple full-thickness serial optical sections using the microscope acquisition software. Typically, to image the entire eye using the 10X objective lens, the microscope defines a square area consisting of 2 x 2 (XY) field-of-view with 10% overlap between them. Using the motorized platform, the microscope automatically acquires the four fields-of-view in a sequential pattern and uses information from the overlapping margins to stitch the individual field-of-view into a single image. Raw image files were imported into ImageJ/Fiji (NIH) using Bio-Formats or to Imaris (Bitplane) for further analysis. For cell counts and quantitative clonal analyses, supervised image segmentation and blob detection was performed on individual optical sections. Identified blobs were manually validated and their number, size and signal intensity as mean grey values were measured.

### Statistical analysis

Sample sizes were not pre-determined, but are similar with what was reported previously (Farrelly et al. 2021). Data were collected and quantified randomly, and their distribution was assumed normal, but this was not formally tested. Lineage tracing, photo-labeling and wounding experiments were successfully reproduced under similar conditions using different mouse cohorts. Unless stated otherwise in the legends, data presented in the figures are from a single cohort of at least three mice, imaged in tandem, using identical experimental parameters. The values of “N” (sample size) refer to data points obtained from all mice within the cohort, unless otherwise indicated, and are provided in the figure legends. Statistical calculations and graphical representation of the data were performed using the Prism software package (GraphPad). Data are expressed as percentages or mean ± S.E.M and unpaired Student’s *t*-test was used to analyze data sets with two groups, unless otherwise stated in the figure legends. For all analyses, *P*-values < 0.05 were designated as significant and symbolized in figure plots as \**P* < 0.05, \*\**P* < 0.01, \*\*\**P* < 0.001, \*\*\*\**P* < 0.0001, with precise values supplied in figure legends. No data were excluded from the analysis.

## Notes

### Competing Interest Statement

The authors have declared no competing interest.

### Summary of Updates

New data: 1) Single-cell transcriptomic analysis of the mouse corneal endothelium during aging (Fig. 1 H-I, Fig. S2). 2) New complementary laser-induced injury model to test corneal endothelial cell proliferation during wound healing. (Fig. 2 E-F, Fig S4 A-H) 3) Histological analysis of the corneal endothelium from corneas injected the pro-regenerative mRNA factors. (Fig 4 I) 4) Long-term follow-up of lineage traced corneal endothelial cells from corneas injected the pro-regenerative mRNA factors. (Fig S10 C) Other revisions: 1) Revised title. 2) Revised abstract. 3) Various edits to the main text and Methods to improve clarity.

